# Single-Cell Multiomic Profiling Uncovers Radiation Dosage-Sensitive, Cluster-Specific Regulatory Dynamics in Glioblastoma

**DOI:** 10.64898/2025.12.01.691699

**Authors:** Khoi Huynh, Sara Nalina Barcik Weissman, Cheol Park, Maria L.G. Quiloan, Blake Sun Chang, Eric Chang, Kelly Street, David Tran, Aram S. Modrek

## Abstract

Glioblastoma (GBM) is an aggressive brain tumor that inevitably recurs after chemoradiotherapy, resulting in poor patient outcomes. Given GBM’s cellular heterogeneity, we hypothesized that radiation induces sub-population specific alterations to survive and adapt to radiation stress. We performed integrated single-cell RNA-seq and ATAC-seq analyses in Glioma Stem-Like Cultures three hours after they were exposed to clinically relevant radiation doses. Radiation reshaped the cellular landscape, altering both cell type composition and GBM subtype distribution. Cluster-specific and shared transcriptional programs were induced in a dose-dependent manner, with differentially expressed genes enriching distinct biological pathways. Chromatin accessibility analyses revealed parallel cluster-specific remodeling, with both opening and closing of regulatory elements linked to functional pathway shifts. Notably, 2 Gy and 6 Gy exposures elicited conserved transcriptional profiles in RNA clusters across different radiation doses. Together, these results highlight immediate radiation-induced transcriptional and chromatin remodeling programs in GBM at single-cell resolution and identify conserved cluster-specific adaptations that may underlie therapeutic resistance.

## INTRODUCTION

Despite the aggressive nature and poor prognosis of glioblastoma (GBM), therapeutic options remain limited and continue to pose a significant clinical challenge. The recurrence of GBM is almost inevitable, even after aggressive multimodal treatment involving surgical resection, radiotherapy, chemotherapy with temozolomide, and tumor-treating fields ^1^. This underscores the urgent need to better understand the biological mechanisms driving tumor relapse. While numerous investigations have explored various contributing factors, such as the roles of genetics and epigenetics in the context of radiation resistance, few studies address how subpopulations of tumor cells respond and adapt acutely (within hours) to radiation stress.

Many genes are individually linked to radiation resistance. However, studies have begun to uncover the diverse cell states within glioblastoma (GBM) and their relationship to therapy response ^2–4^. GBM heterogeneity is influenced not only by intrinsic factors but also varies significantly between newly diagnosed and recurrent tumors, as well as treatment-naïve and previously treated patients ^5^. Understanding complex cellular networks that drive glioblastoma (GBM) heterogeneity and treating the tumor as a population of genetically and epigenetically diverse individual cells may be important for developing more effective therapies ^6–9^.

Given the cellular heterogeneity of glioblastoma (GBM), we believe investigating the tumor at the single-cell level is necessary to uncover distinct gene-expression and chromatin alterations that may drive recurrence and treatment resistance. In this study, we focus on the 3-hour post-irradiation timepoint, which represents a window when cells actively respond to radiation-induced DNA damage but have not yet undergone selection due to apoptosis or necrosis ^10^. This timing allows us to capture early transcriptional and chromatin accessibility changes that reflect the cellular biological response to radiation. We characterize the changes in the landscape of gene expression and chromatin accessibility using scRNA-seq and scATAC-seq to provide insight into the initial single-cell-level molecular events that may influence therapeutic outcomes.

## RESULTS

### Radiation reshapes cell composition and GBM subtype distribution, as revealed by integrated scRNA-seq and scATAC-seq

We performed radiation treatments at three doses: 0 Gy (control), 2 Gy, and 6 Gy. After quality control measures using the criteria described in the method, all samples were combined for clustering analysis. Using the Louvain algorithm in Seurat ^11^, we identified five distinct RNA Clusters (0, 1, 2, 3, and 4) based on transcriptomic profiles across all treatment groups (Figure 1A). Pairwise comparisons of cell proportions per cluster revealed a significant reduction in Cluster 4 cells in the 6 Gy group, dropping to 33.36% compared to 0 Gy (p = 0.000016; Figure 1B and Table S1). Next, we annotated each RNA cluster using established GBM subtype markers as reported by Verhaak and colleagues ^12^. These markers correspond to GBM Classical, Mesenchymal, and Proneural subtypes. Figure 1C illustrates the distribution of GBM subtypes across clusters and treatment conditions. Both 2 Gy and 6 Gy treatments significantly increased the proportion of Classical subtype cells in Clusters 0, 1, and 2 compared to 0 Gy (p < 0.05) within 3 hours of radiation. In Cluster 4, 2 Gy also increased the Classical subtype cell ratio relative to 0 Gy. Conversely, the 0 Gy group had a higher proportion of Mesenchymal subtype cells across all clusters (0 - 4) compared to 2 Gy, and 6 Gy further reduced Mesenchymal cells in Clusters 1 and 4. For the Proneural subtype, 0 Gy showed higher proportions in Clusters 1 and 3 compared to 2 Gy, and in Clusters 1 and 4 compared to 6 Gy. The results suggest that radiation, at both low and high doses, induces a transcriptional shift toward the Classical subtype in glioblastoma stem cells (GSCs) as early as 3 hours post-treatment. At this time point, cell death has not yet occurred, indicating that the observed changes likely reflect early stress responses or survival signaling rather than selective cell loss. This shift is unexpected given that numerous studies have shown the Mesenchymal subtype to be more resistant to radiation and associated with aggressive, therapy-refractory phenotypes ^13–17^.

**Figure 1.**
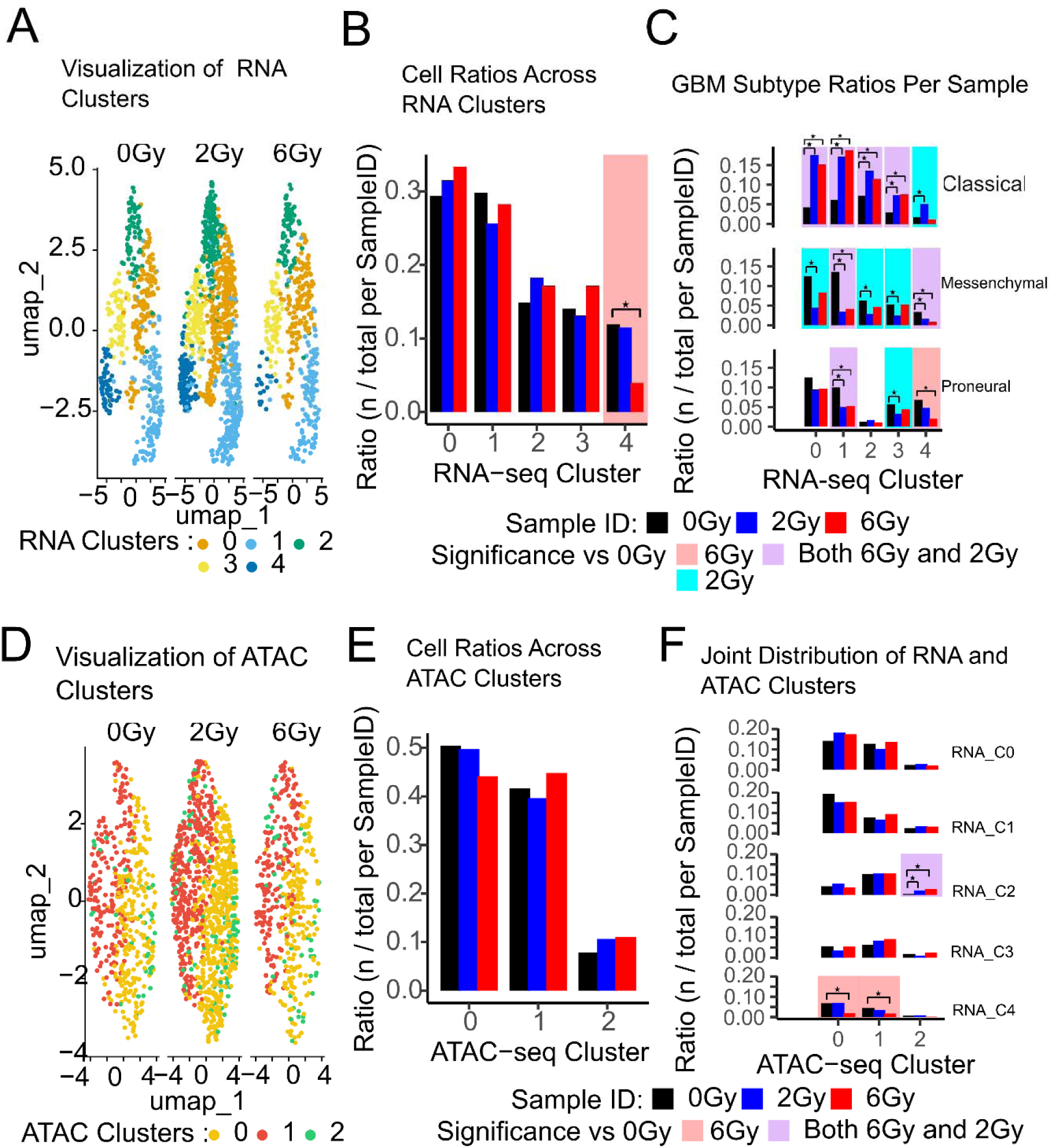
Radiation reshapes cell composition and GBM subtype distribution, as revealed by integrated scRNA-seq and scATAC-seq. (A) UMAP projection of RNA-seq profiles showing transcriptionally defined clusters across all samples. (B) Pairwise statistical comparison of cell ratios across RNA-defined clusters using chi-squared test (p-value□<□0.05). (C) Distribution of GBM molecular subtypes across RNA-defined clusters, with pairwise comparisons performed using chi-squared test with significance defined as p-value < 0.05. (D) UMAP projection of ATAC-seq profiles showing chromatin accessibility-based clustering across all samples. (E) Cell ratio distributions across ATAC-defined clusters were compared between samples using a chi-squared test. Statistical significance was determined at a threshold of *p* < 0.05. (F) Cross-modal comparison of cell ratio distributions per RNA-defined cluster within each ATAC-defined cluster for individual samples, with statistical evaluation performed using chi-squared test. Statistical significance was determined at a threshold of *p* < 0.05.

Several potential explanations may account for this observation. First, enrichment for Classical subtype markers may be due to a rapid transcriptional response to radiation that temporarily dominates the transcriptional signature before other resistance mechanisms emerge. For example, Classical GSCs have been shown to modulate neurodevelopmental pathways and shift toward a quiescent neural stem cell-like state following radiation, which may confer transient survival advantages ^18–21^. Second, radiation may induce a stress-mediated reprogramming that favors Classical gene expression signatures, possibly through p53-dependent pathways or chromatin remodeling events that are more active in Classical cells ^22,23^. Third, the observed shift could reflect a transient phenotypic plasticity rather than a stable subtype transition. GSCs are known to exhibit dynamic interconversion between subtypes in vitro, and radiation may trigger a temporary reversion to a Classical-like state before eventual transition to Mesenchymal features associated with recurrence and resistance ^24,25^.

After clustering cells based on RNA profiles, we examined their grouping by ATAC-seq profiles. Using the same Louvain clustering method in Seurat, cells were grouped into three ATAC clusters (Figure 1D). Surprisingly, pairwise comparisons showed no significant increase in cell proportions within any ATAC cluster for the radiation-treated groups (2 Gy, 6 Gy) compared to the control (0 Gy) (Figure 1E). We then investigated whether cells assigned to specific RNA clusters were evenly distributed across ATAC clusters or showed preferential localization. Cells from RNA cluster 4 in the 6 Gy sample were more frequently found in ATAC Clusters 0 and 1 compared to 0 Gy. Similarly, RNA Cluster 2 cells in the 2 Gy sample were more enriched in ATAC Cluster 2 (Figure 1F). Full statistical details for all pairwise comparisons are provided in Tables S1, S2, S3, and S4.

### Radiation dose–dependent gene expression reveals subtype-defined and overlapping molecular pathways in GBM

To examine transcriptional responses to different radiation doses, we performed pathway enrichment and differential expression analyses across RNA clusters following 2 Gy and 6 Gy treatments (Figure 2). In Figure 2A, we use Venn diagrams to show the numbers of differentially expressed genes in response to 2 Gy and 6 Gy radiation, as well as any overlap. RNA cluster 4 showed notable shifts in cell proportions following radiation treatment; however, it was excluded from further analysis because the number of cells in each condition fell below our minimum threshold of 100 cells required for reliable differential gene expression analysis. To investigate radiation-induced transcriptional changes, we performed differential gene expression analysis within the four largest RNA clusters (0, 1, 2, and 3), comparing 2 Gy and 6 Gy treatments to the control (0 Gy). Overall, the 6 Gy treatment resulted in substantially more differentially expressed genes (DEGs with p-value < 0.05) than 2 Gy, with 6.19-, 9.24-, 5.53-, and 1.57-fold increases in Clusters 0, 1, 2, and 3, respectively (Figure 2A). Additionally, Clusters 0, 1, 2, and 3 shared 618, 524, 333, and 196 DEGs between both treatment groups when compared to the control (Figure 2A). Our findings show that higher radiation dose (6 Gy) induces more transcriptional response across major RNA clusters compared to 2 Gy. The shared DEGs between treatment groups suggest common radiation-responsive pathways, while the magnitude of change highlights dose-dependent effects.

**Figure 2.**
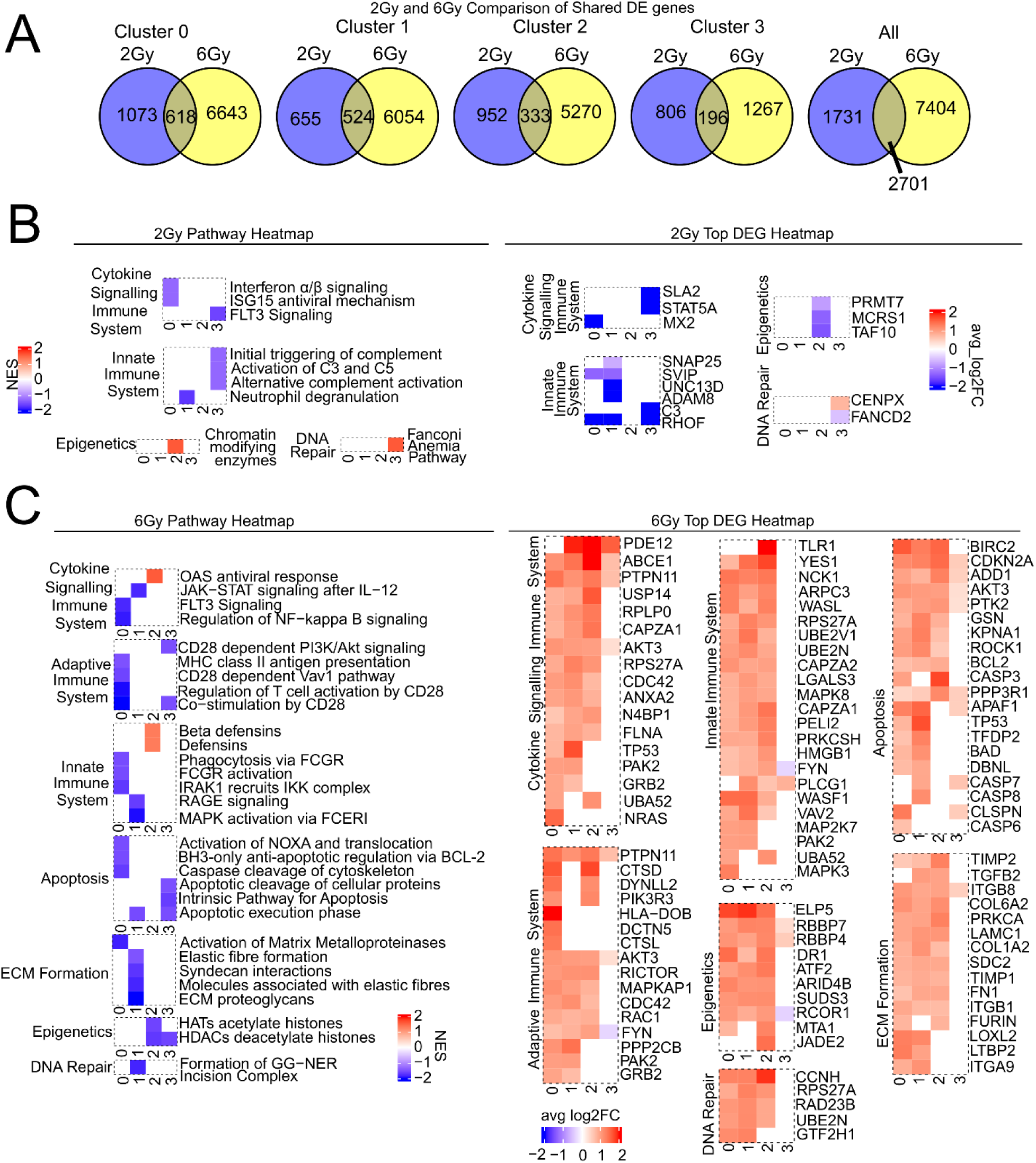
Radiation dose–dependent gene expression reveals subtype-defined and overlapping molecular pathways in GBM. (A) Venn diagrams showing the overlap of differentially expressed genes (DEGs) between 2 Gy and 6 Gy conditions. The left panels visualize overall DEG overlap count per RNA cluster, and the right panel shows overall DEG overlap regardless of RNA cluster. DEGs were identified using |log₂FoldChange| ≥ 0.1 and p-value < 0.05. (B) Left panel: heatmap of normalized enrichment scores for pathways enriched (restricted to DEGs unique to 2 Gy vs. 0 Gy). Statistical significance was determined by GSEA (p-value < 0.05). Right panel: heatmaps visualize DEGs found in enriched pathways from the left panel. For every cluster and pathway, the top five up- and down-regulated DEGs with p-value < 0.05 were selected for visualization. Some pathways lack strongly negative (or positive) log₂FoldChange DEGs associated with enriched GSEA pathways (left panel). (C) Left panel: heatmap of normalized enrichment scores for pathways enriched (restricted to DEGs unique to 6 Gy vs. 0 Gy). Statistical significance was determined by GSEA (p-value < 0.05). Right panel: heatmaps visualize DEGs found in enriched pathways from the left panel. For every cluster and pathway, the top five up- and down-regulated DEGs with p-value < 0.05 were selected for visualization. Some pathways lack strongly negative (or positive) log₂FoldChange DEGs associated with enriched GSEA pathways (left panel).

When we performed a global differential expression analysis independent of RNA cluster identity, the number of shared DEGs between the treatment groups totaled 2701 (“all” venn diagram in Figure 2A). These results suggest that the higher radiation dose (6 Gy) induces stronger gene upregulation, with over 80% of DEGs being upregulated. In contrast, the 2 Gy treatment primarily leads to gene suppression, with more than 80% of DEGs being downregulated (Figure S1A). This directional shift in gene expression is consistent with previously published findings^26^. To further characterize these changes, we categorized DEGs into common (shared between 2 Gy and 6 Gy) and unique (specific to either 2 Gy or 6 Gy) groups. A comprehensive list of all DEGs can be found in Data S1.

For the pathways heatmap and DEGs heatmap in Figure 2B and 2C, we showed gene expression and pathway analysis results for DEGs that are unique to either 2 Gy or 6 Gy radiation response. Furthermore, we focused on immune-related pathways, epigenetic regulation, apoptosis, extracellular matrix (ECM) formation, and DNA repair because these processes are well-documented as central components of cellular responses to ionizing radiation. Each pathway plays a critical role in determining radiation sensitivity or resistance through mechanisms such as immune modulation, chromatin remodeling, programmed cell death, tissue remodeling, and repair of radiation-induced DNA damage.

Pathway analysis on 2 Gy only DEGs revealed selective activation of immune-related and epigenetic processes across clusters. Cytokine signaling pathways, including interferon-α/β signaling, ISG15 antiviral mechanism, and FLT3 signaling, were suppressed in Clusters 0 and 3, along with the suppressed innate immune mechanisms such as complement activation and neutrophil degranulation in Clusters 1 and 3 (2 Gy pathway heatmap in Figure 2B). Epigenetic regulation through chromatin-modifying enzymes and DNA repair via the Fanconi Anemia pathway were also enriched in Clusters 2 and 3 (Figure 2B), respectively. Examination of top differentially expressed genes (DEGs) showed downregulation of SLA2 (p-value = 0.015, log_2_FC = -2.97 in RNA Cluster 3), STAT5A (p-value = 0.048, log_2_FC = -2.97 in RNA Cluster 3), and MX2 (p-value = 0.00077, log_2_FC = -3.75 in RNA Cluster 0) in cytokine signaling pathways, while PRMT7 (p-value = 0.033, log_2_FC = -0.84 in RNA Cluster 2) and MCRS1 (p-value = 0.044, log_2_FC = - 1.36 in RNA Cluster 2) were notably suppressed within the chromatin-modifying enzyme pathway (Figure 2B). In the Fanconi Anemia pathway, CENPX (p-value = 0.034, log_2_FC = 0.76 in RNA Cluster 3) was upregulated, whereas FANCD2 (p-value = 0.035, log_2_FC = -0.53 in RNA Cluster 3) was downregulated (Figure 2B), suggesting early engagement of DNA damage recognition components rather than full repair activation at this time point.

Cells exposed to 6 Gy exhibited broader and more pronounced unique transcriptional changes, with clear cluster-specific patterns. Pathway enrichment indicated activation of the OAS antiviral response and JAK–STAT signaling in Clusters 2, alongside innate immune system pathways, such as β-defensins and defensins. Beyond differences in immune-related pathways, RNA clusters exhibited distinct transcriptional responses in other pathways to 6 Gy radiation (Figure 2C). Cluster 1 showed strong suppression of pathways related to extracellular matrix (ECM) organization, including ECM proteoglycans, elastic fiber formation, and Syndecan interactions. Genes such as ITGB1 (p-value = 6.60 x 10^-8^, log_2_FC = 0.72 in RNA Cluster 1) and FN1 (p-value = 1.08 x 10^-8^, log_2_FC = 0.91 in RNA Cluster 1) were upregulated and involved in all three suppressed ECM-related pathways (Figure 2C), suggesting structural remodeling under high-dose stress.

Clusters 0 and 3 displayed downregulation of multiple apoptosis-related pathways, including activation of NOXA and BCL-2 regulation, indicating a transcriptional pattern that may reduce programmed cell death (Figure 2C). Cluster 2 remained distinct, with radiation primarily affecting immune-related pathways and epigenetic regulation. Both histone acetyltransferase (HAT) and histone deacetylase (HDAC) pathways were suppressed (Figure 2C), despite their typically opposing roles ^27^. This concurrent downregulation implies a broader reduction in histone modification activity rather than a shift in the balance between acetylation and deacetylation. Since histone modifications are essential for maintaining dynamic chromatin states, their suppression may result in a more static chromatin configuration. A less flexible chromatin environment can hinder the accessibility and recruitment of DNA repair machinery, as supported by previous studies showing that reduced histone modification activity compromises DNA damage response efficiency ^28–30^. Therefore, while our data captures an early transcriptional response, these changes may predispose cells to compromised DNA damage repair at later stages.

Together, these findings indicate dose-dependent and cluster-specific transcriptional responses to radiation. At 2 Gy, clusters primarily activate immune signaling and initiate DNA repair, whereas at 6 Gy, clusters broadly suppress apoptosis and epigenetic factors while enhancing antiviral and immune pathways. These patterns suggest distinct adaptive responses that may influence survival and therapeutic resistance.

### Shared DEGs reveal distinct functional pathways and both common and cluster-specific transcriptional responses in GBM

To characterize potentially shared transcriptional responses to radiation, we conducted differential gene expression and pathway enrichment analyses across RNA clusters following exposure to 2 Gy and 6 Gy (Figure 3A,3B). Figures 3A and 3B present results for genes that are differentially expressed (p-value < 0.05) in response to both radiation doses, allowing for the identification of shared transcriptional radiation response programs. We visualized immune signaling, epigenetic regulation, apoptosis, extracellular matrix organization, and DNA repair processes. These pathways are recognized as mediators of radiation-induced cellular effects and are involved in modulating radiation sensitivity through mechanisms such as immune activation or suppression, chromatin remodeling, programmed cell death, structural tissue remodeling, and the repair of DNA lesions.

**Figure 3.**
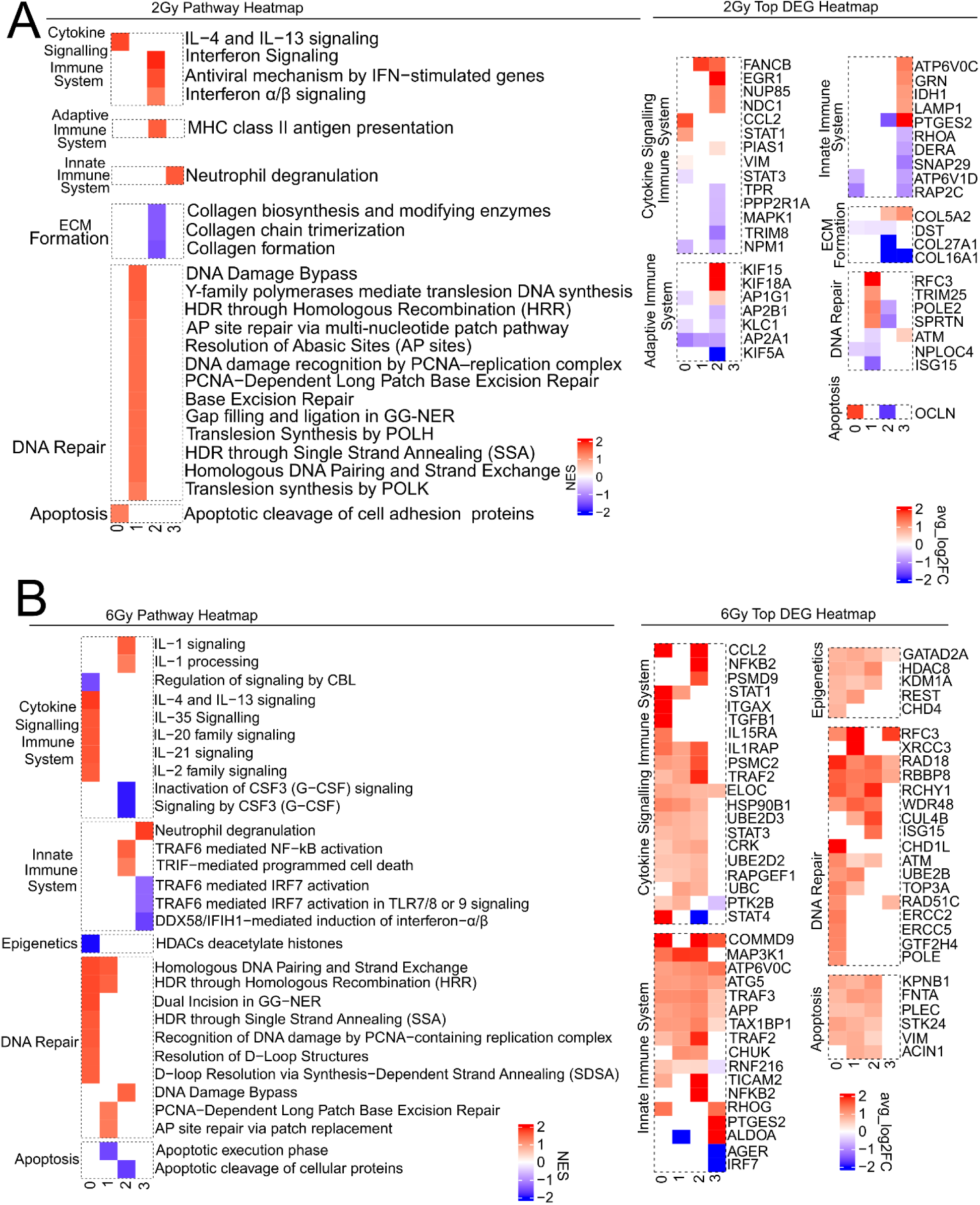
Shared DEGs reveal distinct functional pathways and both common and cluster-specific transcriptional responses in GBM. (A) Left panel: heatmap of normalized enrichment scores for pathways enriched (restricted to shared DEGs (found in response to both 2 Gy and 6 Gy) to 2 Gy vs. 0 Gy). Statistical significance was determined by GSEA (p-value < 0.05). Right panel: heatmaps visualize DEGs found in enriched pathways from the left panel. For every cluster and pathway, the top five up- and down-regulated DEGs with p-value < 0.05 were selected for visualization. Some pathways lack strongly negative (or positive) log₂FoldChange DEGs associated with enriched GSEA pathways (left panel). (B) Left panel: heatmap of normalized enrichment scores for pathways enriched (restricted to shared DEGs (found in response to both 2 Gy and 6 Gy) to 6 Gy vs. 0 Gy). Statistical significance was determined by GSEA (p-value < 0.05). Right panel: heatmaps visualize DEGs found in enriched pathways from the left panel. For every cluster and pathway, the top five up- and down-regulated DEGs with p-value < 0.05 were selected for visualization. Some pathways lack strongly negative (or positive) log₂FoldChange DEGs associated with enriched GSEA pathways (left panel).

RNA Cluster 0 showed enrichment of multiple interleukin (IL) signaling pathways, including IL-4, IL-13, IL-35, IL-20, IL-21, and IL-2. IL-4 and IL-13 signaling pathways were activated under both 2 Gy and 6 Gy radiation conditions. This activation may be driven by the consistent upregulation of CCL2 and STAT1 in both treatment groups, suggesting a regulatory role in initiating the IL-4/IL-13-mediated transcriptional response to radiation. Despite other shared DEGs between 2 Gy and 6 Gy (Figure 3A,3B), IL-35, IL-20, IL-21, and IL-2 are uniquely upregulated in response to 6 Gy radiation. Cluster 0 also exhibited activation of apoptosis-related pathways under 2 Gy (Figure 3A) and enrichment of DNA repair pathways under 6 Gy (Figure 3B), including homologous recombination (HRR), single-strand annealing (SSA), PCNA-dependent damage recognition, and global genome nucleotide excision repair (GG-NER), indicating diverse responses to DNA damage among subpopulations.

RNA Cluster 1 did not show enrichment or suppression of immune-related pathways but exhibited DNA repair pathway activation under both radiation doses. These included HRR, PCNA-dependent long patch base excision repair (BER), and resolution of abasic sites via BER (Figure 3A,3B). Under 2 Gy, GG-NER and SSA were uniquely enriched (Figure 3A), suggesting a dose-specific response to helix-distorting lesions and single-strand breaks. RNA Cluster 2 showed IL-1 signaling enrichment under 6 Gy via shared DEGs (Figure 3B). Granulocyte colony-stimulating factor (G-CSF), which plays a regulatory role in key signaling pathways such as JAK/STAT, MAPK, and PI3K/AKT, was found to be suppressed following radiation. Since G-CSF is essential for promoting the proliferation and differentiation of neutrophil precursors ^31^, its downregulation may reflect a reduced capacity for neutrophil production or mobilization in response to radiation-induced stress. Under 2 Gy, interferon alpha/beta signaling was enriched, along with MHC class I antigen presentation (Figure 3A). These pathways are associated with STING-mediated immune responses ^32,33^. RNA Cluster 3 exhibited suppression of IFN-α and IFN-β production under 6 Gy via downregulation of the TRAF6-dependent IRF7 activation pathway ^34^ (Figure 3B).

In summary, shared DEGs between 2 Gy and 6 Gy radiation treatments revealed distinct transcriptional responses across RNA clusters. Cluster 0 exhibited dose-dependent activation of interleukin signaling and DNA repair pathways. Cluster 1 primarily engaged DNA repair mechanisms without immune pathway involvement. Cluster 2 showed differential regulation of immune signaling, ECM organization, and DNA damage responses depending on radiation dose. Cluster 3 demonstrated immune suppression and enrichment of neutrophil degranulation, with limited additional pathway activity. These results highlight the heterogeneity of radiation-induced transcriptional programs and suggest cluster-specific mechanisms of cellular adaptation.

### Chromatin opening reveals distinct functional pathways and cluster-specific chromatin responses in GBM following radiation

In addition to analyzing gene expression, we investigated changes in chromatin accessibility in response to different radiation doses (2 Gy and 6 Gy) across RNA clusters. To do this, we performed single-cell ATAC-seq (scATAC-seq) followed by differential chromatin accessibility analysis. Then, we performed motif analysis and gene ontology on unique differential chromatin accessibility regions (DCARs) in response to either 2 Gy or 6 Gy radiation. Figure 4A shows the count of DCARs in response to either radiation dose or any overlap for each RNA cluster. The “all” Venn diagram shows the count and overlapped DCARs regardless of RNA clusters. Figure 4B and 4C show the normalized enrichment score for pathway analysis and coverage fold change heatmap for unique DCARs.

**Figure 4.**
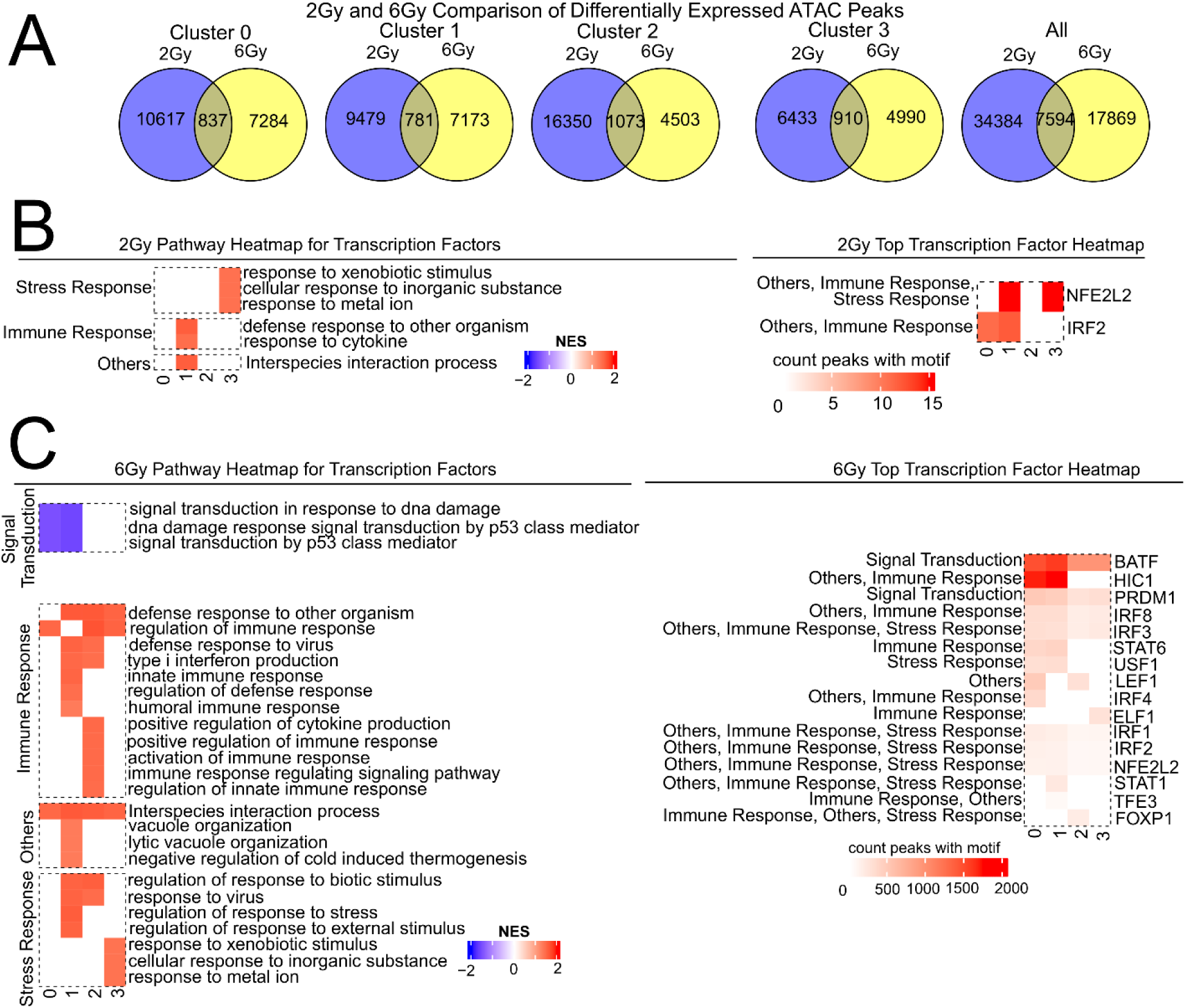
Chromatin opening reveals distinct functional pathways and cluster-specific chromatin responses in GBM following radiation. (A) Venn diagrams showing the overlap of differential chromatin accessibility regions (DCARs) between 2 Gy and 6 Gy conditions. The left panels show cluster-specific overlaps, and the right panel shows the overall DCARs overlap regardless of RNA cluster. DCARs were defined using |log₂ fold change| ≥ 1 and p-value < 0.05. (B) Left panel: heatmap of normalized enrichment scores for pathways enriched (restricted to transcription factors (TFs) binding to open DCARs in response to 2 Gy vs. 0 Gy). Statistical significance was determined by GSEA (p-value < 0.05). Right panel: heatmaps visualize the count of open DCARs (p-value < 0.05 and log2FoldChange > 1) in response to 2 Gy for all TFs found in enriched pathways from the left panel. (C) Left panel: heatmap of normalized enrichment scores for pathways enriched (restricted to transcription factors (TFs) binding to open DCARs in response to 6 Gy vs. 0 Gy). Statistical significance was determined by GSEA (p-value < 0.05). Right panel: heatmaps visualize the count of open DCARs (p-value < 0.05 and log2FoldChange > 1) in response to 6 Gy for all TFs found in enriched pathways from the left panel.

Overall, 2 Gy radiation resulted in greater chromatin accessibility alterations compared to 6 Gy, with 1.46-, 1.32-, 1.29-, and 1.92-fold increases in differential chromatin accessibility regions (DCARs) in RNA Clusters 0, 1, 2, and 3, respectively. Interestingly, RNA Clusters 0, 1, 2, and 3 contained 837, 781, 1073, and 910 DCARs, respectively, that were shared between both radiation treatments (2 Gy and 6 Gy) when compared to the control group (0 Gy). When we analyzed DCARs across all clusters, without considering transcriptional identity (RNA clusters), we identified 7594 regions that were commonly accessible in both radiation conditions relative to the control. Additionally, our analysis showed that 2 Gy treatment tends to close more chromatin regions, while 6 Gy treatment tends to open more (Figure S1B). The magnitude of these changes ranged from a 2-fold to a 125-fold difference. This finding is notable because it suggests that high-dose radiation may initially increase chromatin accessibility within the first few hours after exposure, likely to facilitate the DNA damage response by enabling access for repair machinery. This transient chromatin relaxation is consistent with previous studies showing that ionizing radiation induces rapid chromatin remodeling to allow recruitment of DNA repair proteins such as γH2AX and 53BP1 ^35–38^. However, by 24 hours post-treatment, the observed decrease in accessibility may reflect chromatin recompaction and transcriptional repression, a pattern also reported in ATAC-seq studies of irradiated tissues where reduced accessibility at transcriptional start sites was detected following acute high-dose exposure. These dynamic changes are likely mediated by epigenetic modifications, including histone deacetylation and methylation, which have been shown to regulate chromatin structure during the DNA damage response ^39–41^. A complete list of DCARs can be found on Data S2. To further understand the biological implications of these changes, we identified transcription factors (TFs) that bind to regions that became accessible following radiation exposure. We focused on TFs associated with positively regulated DCARs and performed gene ontology analysis using the GO database to explore their biological functions.

In response to 2 Gy radiation, RNA Cluster 3 showed enrichment of stress response pathways, including responses to xenobiotic stimuli, inorganic substances, and metal ions (Figure 4B). RNA Cluster 1 exhibited enrichment of immune-related pathways such as defense response to other organisms and cytokine signaling, whereas Clusters 0 and 2 did not show significant pathway enrichment. Motif analysis identified TFs associated with these pathways, including NFE2L2 (p-value = 1 x 10^-5^, count of DCARs with motif = 15 in RNA Cluster 3) and IRF2 (p-value = 1 x 10^-9^, count of DCARs with motif = 159 in RNA Cluster 1), which were among the most enriched motifs in 2 Gy radiation-induced open chromatin regions (Figure 4B).

Pathway enrichment was broader in response to 6 Gy and included alterations in signal transduction in response to DNA damage, p53-mediated signaling DNA damage pathways (Clusters 0 and 1), and multiple immune-related processes such as regulation of innate immune responses (Cluster 1 and 2), cytokine production (Cluster 2), and interferon signaling (Cluster 1) (Figure 4C). Stress-related pathways, including responses to xenobiotic stimulus and metal ions, were also enriched in Clusters 1,2, and 3.

Motif analysis revealed a diverse set of TFs, including BATF (p-value = 1 x 10^-461^ and 1 x 10^-513^ with count of DCARs with motif = 1471 and 1587 for RNA Cluster 0 and 1 respectively), IRF family members (IRF1, IRF2, IRF3, IRF8 with p-value < 0.05 and at least 11 associated DCARs in each clusters), STAT1(p-value =1 x 10^-2^ with count of DCARs with motif = 210 for RNA Cluster 1), and NFE2L2 (p-value < 0.05 with average count of DCARs with motif across 4 clusters = 97), indicating activation of immune and stress response programs alongside DNA damage signaling (Figure 4C).

Across clusters, RNA Cluster 3 showed enrichment of stress response pathways under both radiation doses, while immune-related pathways were more prominent after 6 Gy (Figure 4B,4C). RNA Cluster 1 exhibited stress and immune pathway enrichment only after 6 Gy, and Cluster 2 showed enrichment of both immune and stress responses following 6 Gy (Figure 4B,4C). Cluster 0 displayed activation of immune regulation pathways under 6 Gy (Figure 4B,4C). Additionally, 6 Gy suppressed several p53-related DNA damage response pathways in Clusters 0 and 1, coinciding with increased chromatin accessibility at BATF binding sites (Figure 4B,4C).

In summary, our findings reveal that radiation dose influences chromatin accessibility and transcription factor activity in a cluster-specific manner. Low-dose radiation (2 Gy) tends to activate stress responses, while high-dose radiation (6 Gy) induces broader immune responses and alters DNA damage repair pathways, particularly those regulated by p53. Transcription factors such as NFE2L2 and BATF appear to play key roles in mediating these effects, suggesting potential targets for enhancing radiation sensitivity or resistance.

### Differential accessibility of chromatin regions reveals distinct functional pathways and cluster-specific chromatin responses in GBM

To investigate transcriptional programs potentially affected by radiation-induced closure of chromatin regions, we analyzed differentially accessible chromatin regions (DACRs) showing decreased accessibility after 2 Gy and 6 Gy treatments. Motif enrichment was performed to identify transcription factors (TFs) associated with these closed regions, and pathway analysis was conducted for genes containing these motifs (Figure 5A-5D).

**Figure 5.**
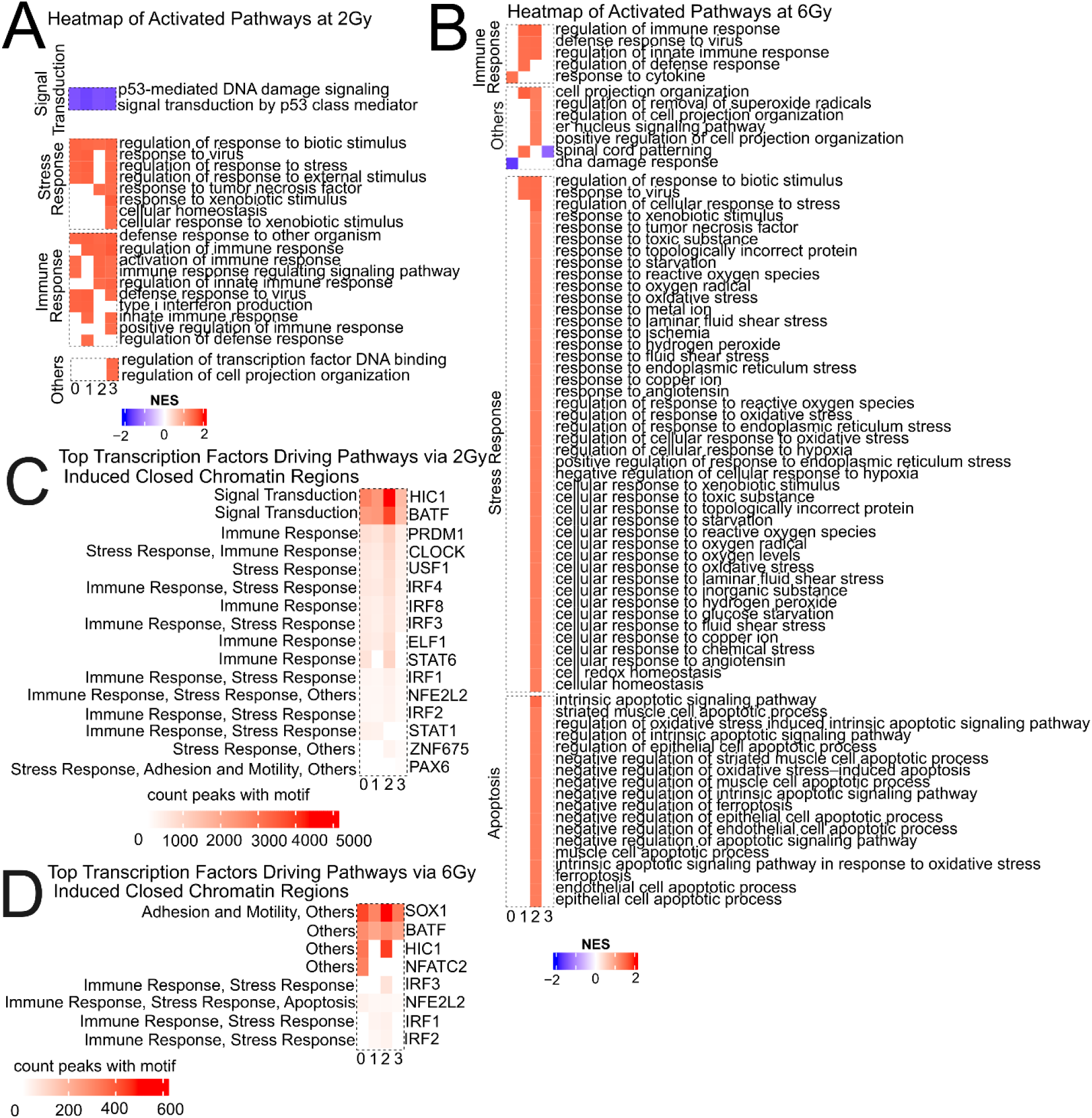
Closure of chromatin regions reveals distinct functional pathways and cluster-specific chromatin responses in GBM. (A) Heatmap of normalized enrichment scores for pathways enriched (restricted to transcription factors (TFs) binding to closed DCARs (p-value < 0.05 and log2FoldChange < -1 in response to 2 Gy vs. 0 Gy). Statistical significance was determined by GSEA (p-value < 0.05). (B) Heatmap of normalized enrichment scores for pathways enriched (restricted to transcription factors (TFs) binding to closed DCARs (p-value < 0.05 and log2FoldChange < -1 in response to 6 Gy vs. 0 Gy). Statistical significance was determined by GSEA (p-value < 0.05). (C) Heatmaps visualize the count of closed DCARs (p-value < 0.05 and log2FoldChange < -1) in response to 6 Gy vs. 0 Gy for all TFs found in enriched pathways from A. (D) Heatmaps visualize the count of closed DCARs (p-value < 0.05 and log2FoldChange < -1) in response to 6 Gy vs. 0 Gy for all TFs found in enriched pathways from the B.

Radiation response pathways are strongly associated with closed chromatin regions, including p53-mediated DNA damage signaling and signal transduction by p53 class mediators, as indicated by negative enrichment scores (Figure 5A). This suggests that these pathways are less represented among closed regions and may remain relatively active. In contrast, pathways shown in red, such as regulation of response to biotic stimulus, immune response regulation, and stress response pathways, were highly enriched among closed regions (Figure 5A), indicating functional suppression due to reduced TF binding. The top TFs ranked by the number of differentially accessible ATAC-seq peaks (DACRs) associated with these closed regions included BATF (p-value < 0.05 and average count of DCARs with motif = 1016 across all clusters), HIC1 (p-value < 0.05 and average count of DCARs with motif = 1141 across all clusters), and IRF family members (IRF1, IRF2, IRF3, IRF4, IRF8), along with STAT6 (p-value < 0.05 and average count of DCARs with motif = 251 across all clusters), NFE2L2 (p-value < 0.05 and average count of DCARs with motif = 80 across all clusters), and others involved in immune and stress regulation (Figure 5C). BATF exhibited the highest number of closed binding sites (>1500), suggesting strong suppression of BATF-driven pathways under 2 Gy.

Chromatin closure following 6 Gy treatment affected a broader set of pathways compared to 2 Gy (Figure 5B). Pathways enriched in closed regions included immune response regulation, cytokine production, response to oxidative stress, and intrinsic apoptotic signaling, indicating suppression of these processes. Pathways with negative enrichment scores, such as p53-mediated signaling, were less represented among closed regions, suggesting retention of activity. The top TFs ranked by the number of differentially accessible ATAC-seq peaks (DACRs) associated with closed regions included SOX1, BATF, HIC1, NFE2L2, and IRF family members (IRF1, IRF2, IRF3) (average p-value < 0.05 and average count of associated DCARs = 560 across all of those TFs and across all clusters), indicating that TFs involved in immune and stress regulation were broadly affected (Figure 5D).

### Conserved transcriptional profiles in RNA clusters associated with radiation treatment

To validate our findings, we compared our data with previously published results from two IDH-wildtype patient-derived glioblastoma cell lines, HF2354 and HF3016, by Johnson in 2021 ^42^. These cell lines were treated with a daily dose of 2.5Gy over four days, followed by a five-day recovery period ^42^. Initially, we compared DEGs between radiation-treated and control samples from our cell line against Johnson’s cell lines without considering cluster identity. Surprisingly, the number of correlated DEGs between our data and Johnson’s Cluster 1 was significantly reduced, although the overall trends remained consistent (Figure S2A). Furthermore, our DEGs in response to 2 Gy did not show significant correlation with HF2354 DEGs (p-value > 0.05). Among correlated DEGs, we found only 1 and 74 overlapping DEGs between the two cell lines in response to 2 Gy and 6 Gy, respectively (Figure S2A). The log_2_FoldChange heatmaps for these overlapping genes are shown in Supplemental Figure 3B. Given the heterogeneity of glioblastoma cell lines, we hypothesized that differences in cellular composition might be obscuring meaningful comparisons. To address this, we projected our RNA-seq clusters onto Johnson’s datasets and found that approximately 90 percent of cells from both HF3016 and HF2354 mapped to our Cluster 1 (Figure 6A). This strong overlap suggested that Cluster 1 represents a shared transcriptional state across datasets. To improve the biological relevance and accuracy of our validation, we focused exclusively on Cluster 1 cells from both our dataset and Johnson’s. This approach allowed us to control cellular heterogeneity and ensure that comparisons were made between transcriptionally similar populations.

**Figure 6.**
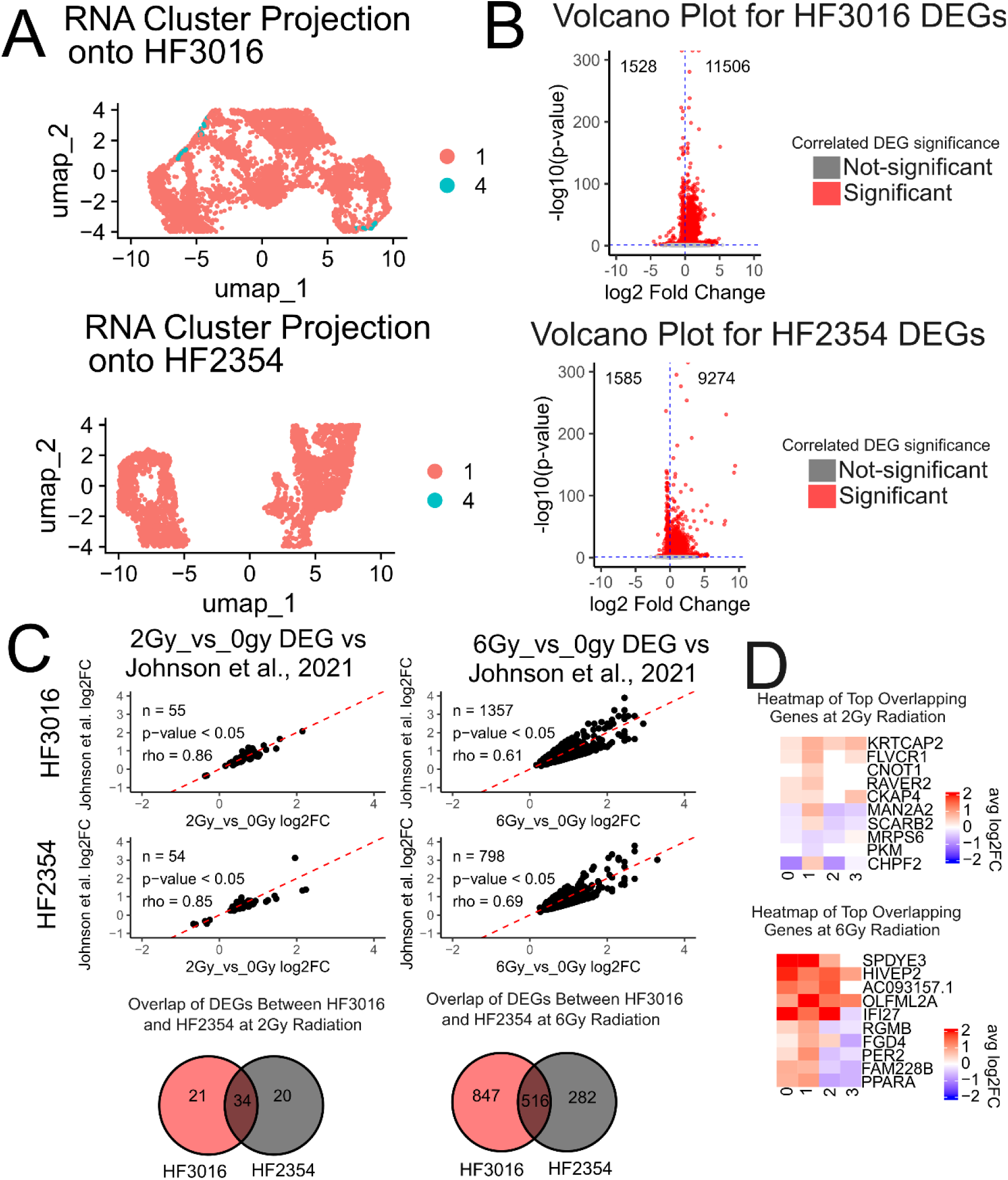
Conserved transcriptional profiles in RNA cluster 1 associated with radiation treatment. (A) RNA projection of our RNA clusters onto Johnson et al., 2021 HF2354 and HF3016 cell lines using dimensionality reduction for cross-sample alignment with Seurat. (B) Volcano plots showing differentially expressed genes (DEGs) identified in Johnson et al., 2021 HF2354 and HF3016 cell lines by comparing treatment groups (2 Gy or 6 Gy) to their respective controls. Each point represents a gene, with log₂FoldChange on the x-axis and –log₁₀ p-value on the y-axis. Genes with p-values < 0.05 are highlighted in red. (C) Volcano plots showing correlated DEGs between our GSC20 and Johnson et al., 2021 HF2354 and HF3016 cell lines. Bottom Venn diagrams illustrate the overlap of correlated DEGs between our GSC20 and HF2354 / HF3016 cell lines in response to 2 Gy (left) and 6 Gy (right). Correlation was assessed using Pearson correlation (p-value < 0.05) combined with differential expression criteria (p-value < 0.05). (D) Heatmaps display log2FoldChange values for the top five up- and down-regulated overlapping genes identified in panel C in response to 2 Gy (top) and to 6 Gy (bottom). Genes and samples were clustered using Euclidean distance, and differential expression was assessed with DESeq2 (p < 0.05).

Differential gene expression analysis of Cluster 1 cells from HF3016 and HF2354 identified 13035 and 10860 differentially expressed genes (DEGs), respectively (p-value < 0.05), with the majority showing increased expression relative to their corresponding controls (Figure 6B). To facilitate a comprehensive comparison across datasets, we included all statistically significant genes without applying a fold-change cutoff. To evaluate the similarity between our data and Johnson’s dataset, we compared DEGs from our Cluster 1 cells to those from Johnson’s Cluster 1. We retained genes with log2FoldChange values that were directionally consistent and within 60% of each other. Using these criteria, we identified 55 and 1357 correlated DEGs between our 2 Gy condition and Johnson’s HF3016 and HF2354 datasets, respectively, and 54 and 798 correlated DEGs between our 6 Gy condition and the same datasets. The corresponding correlation coefficients (ρ) were 0.86, 0.61, 0.85, and 0.69 (Figure 6C). Notably, we found 34 and 516 DEGs shared between the two cell lines in response to 2 Gy and 6 Gy treatments, respectively (Figure 6C).

A heatmap of the top five and bottom five DEGs based on average log_2_FoldChange among the overlapping genes revealed two particularly interesting candidates (Figure 6D). The first is PKM, a gene correlated with our 2 Gy condition. PKM has previously been associated with high radiation resistance, poor patient survival, increased proliferation and migration, and reduced apoptosis in U87 MG cells ^43^. PKM is downregulated in response to 2 Gy radiation in our dataset and similarly downregulated in Johnson’s dataset following fractionated 10 Gy radiation. A second gene of interest is PER2, whose expression is similarly regulated in both our 6 Gy condition and Johnson’s dataset. PER2 has been shown to promote radiation resistance by enhancing DNA repair and mitochondrial function ^44^. In both our data and Johnson’s, PER2 is upregulated in response to radiation, suggesting a role in glioma’s radiation response.

In summary, our comparative analysis with previously published datasets supports the robustness of our findings. The consistent clustering of HF3016 and HF2354 cells into our RNA Cluster 1, along with the correlation of key DEGs such as PKM and PER2, highlights shared transcriptional responses to radiation in stem-like glioblastoma models.

### Enriched and context-dependent pathways emerge from dose-correlated DEGs, revealing a conserved expression signature overlapping with Johnson et al. RNA clusters

To investigate biological processes associated with correlated DEGs, we performed Gene Ontology analysis using Johnson’s fractionated radiation DEGs for HF3016 and HF2354 cell lines and filtered results based on six key functional categories: DNA repair, immune system, extracellular matrix (ECM) formation, cell cycle, cellular response to stress, and epigenetics (Figure 7A–C).

**Figure 7.**
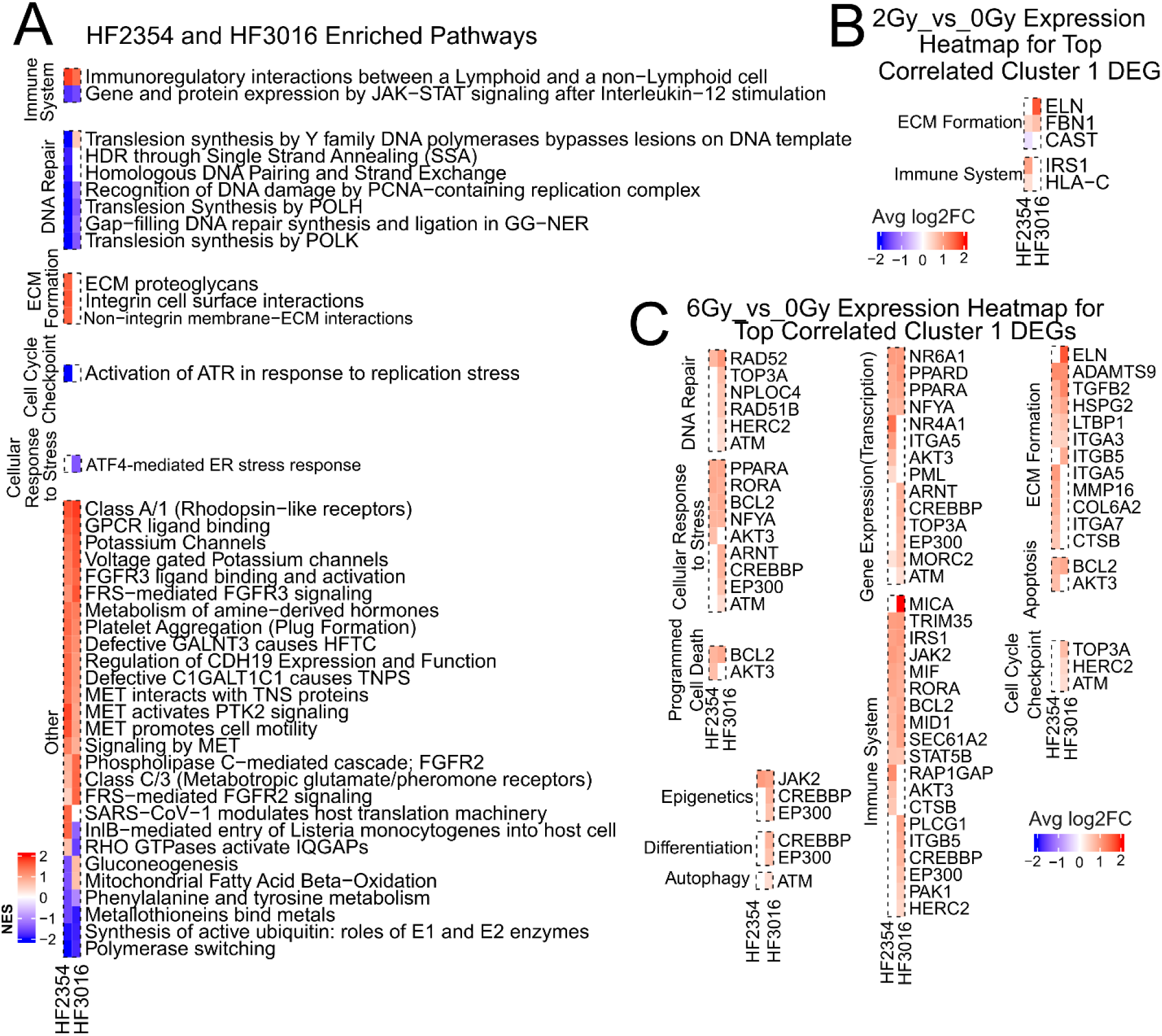
Enriched and context-dependent pathways emerge from dose-correlated DEGs, revealing a conserved expression signature overlapping with Johnson’s Cluster 1. (A) Heatmaps of normalized enrichment scores (NES) for pathways enriched by differentially expressed genes (DEGs) reported by Johnson et al. in HF3016 cells and HF2354 cells. Pathway enrichment was assessed using gene set enrichment analysis (GSEA), with significance defined as p-value < 0.05. (B) Gene expression heatmaps for the correlated DEGs associated with the pathways in A in response to 2 Gy (p < 0.05). For every cluster and pathway, the top five up- and down-regulated DEGs were selected; Some pathways lack strongly negative (or positive) log₂FoldChange DEGs associated with enriched GSEA pathways. (C) Gene expression heatmaps for the correlated DEGs associated with the pathways in A in response to 6 Gy (p < 0.05). For every cluster and pathway, the top five up- and down-regulated DEGs were selected; Some pathways lack strongly negative (or positive) log₂FoldChange DEGs associated with enriched GSEA pathways.

In HF3016 cells, suppressed pathways included DNA damage recognition by PCNA complex, translesion synthesis by POLH and POLK, and immune-related processes such as IL-12 mediated JAK-STAT signaling (Figure 7A). Additionally, pathways involved ATF4-mediated ER stress response, polymerase switching, and ubiquitin-synthesis regulation are also suppressed (Figure 7A). HF2354 cells exhibited similar immune pathway suppression, along with DNA repair processes such as homologous DNA repair via single-strand annealing (SSA) and global genome nucleotide excision repair (GG-NER) (Figure 7A). ECM-related pathways were commonly enriched in both HF2354 and HF3016, including ECM proteoglycan interactions and non-integrin membrane ECM interactions, as well as signaling pathways such as MET receptor signaling and GPCR ligand binding (Figure 7A).

In the 2 Gy treatment arm, correlated DEGs were primarily associated with ECM formation and immune system pathways (Figure 7B). HF3016-correlated DEGs included ECM formation genes (ELN (p-value = 0.048, log_2_FC = 1.57), FBN1 (p-value = 0.049, log_2_FC = 0.55)), while HF2354-correlated DEGs included ECM formation genes (FBN1 (p-value = 0.049, log_2_FC = 0.55), CAST (p-value = 0.0055, log_2_FC = -0.25)) in addition to immune system genes IRS1 (p-value = 0.048, log_2_FC = 0.57) and HLA-C (p-value = 0.0025, log_2_FC = 0.52), indicating shared involvement in ECM organization and immune regulation.

Heatmaps of DEGs correlated with fractionated datasets revealed distinct patterns for HF3016 and HF2354 correlation to 6 Gy treatment response (Figure 7C). HF3016-correlated DEGs included genes involved in DNA repair such as TOP3A (p-value = 0.032, log_2_FC =0.68), ATM (p-value = 0.016, log_2_FC =0.44), HERC2 (p-value = 0.00044, log_2_FC = 0.56), and ECM formation genes such as ITGB5 (p-value = 0.00023, log_2_FC = 1.00). HF2354-correlated DEGs included transcription-related gene expression gene NR4A1 (p-value = 0.023, log_2_FC = 2.13), and ECM-associated gene ITGA5 (p-value = 0.03, log_2_FC = 0.76) (Figure 7C).

## DISCUSSION

We treated glioblastoma cells with 2 Gy and 6 Gy doses of radiation and profiled transcriptomic and epigenetic responses using scRNA-seq and scATAC-seq. RNA clustering revealed dose-dependent shifts in cell proportions across RNA clusters and across GBM subtypes. Differential gene expression and gene set enrichment analyses identified radiation-affected pathways, with results validated against external datasets. ATAC clustering showed stable subpopulation proportions across ATAC clusters, but differential accessibility and motif enrichment analyses revealed transcription factor activity changes linked to RNA-defined clusters. Notably, RNA clusters are localized within specific ATAC clusters post-radiation, suggesting coordinated transcriptional and chromatin remodeling responses that shape cell identity in a dose-specific manner.

Our transcriptomic analysis of glioblastoma stem-like cells (GSCs) exposed to 2 Gy and 6 Gy radiation reveals highly heterogeneous, cluster-specific responses shaped by dose intensity. Pathway signaling and gene expression analyses showed that both doses induce adaptive responses that may select for resistance and recurrence. At 2 Gy, clusters exhibited enrichment of DNA repair pathways such as homologous recombination, base excision repair, and translesion synthesis, with upregulation of FANCB and POLE2, indicating activation of sublethal repair processes ^45,46^. Because FANCB is upregulated and FANCD2 is downregulated, the cells may not fully repair DNA damage and instead rely on PI3K/AKT signaling (EGR1, GRN) ^47,48^, allowing survival under sublethal stress. Simultaneous suppression of cytokine, interferon, and complement signaling genes (MX2, ISG15, STAT5A) could promote immune evasion ^49–51^. Enrichment of DNA repair, chromatin-modifying enzymes, ECM formation (VIM, COL5A2), and Fanconi anemia-related genes may allow repair-competent cells to survive sublethal damage and contribute to tumor repopulation ^52,53^.

In contrast to 2 Gy, which is considered standard fractionation, exposure to 6 Gy irradiation (considered high-dose radiation), downregulated apoptotic and immune signaling pathways (NF-κB, JAK-STAT, CD28 co-stimulation) as well as chromatin remodeling, while upregulating pro-survival and oncogenic drivers (AKT3, BIRC2, NRAS) alongside interleukin signaling (IL-2, IL-21, IL-35). These changes may favor the selection of radioresistant, mesenchymal-like, stem-like populations with enhanced DNA repair capacity and anti-apoptotic properties ^54^. Shared activation of IL-2, IL-4, IL-13, and IL-21 suggests convergent activation of STAT3 and PI3K/AKT pathways that contribute to tumor survival and immune suppression ^54^. Cluster-level analysis further revealed that after 2 Gy, Cluster 1 showed DNA repair and chromatin remodeling gene upregulation (FANCB, RFC3, POLE2, KAT2A) ^55^, while Cluster 0 showed immune suppression and Fanconi anemia pathway activation. After 6 Gy, Cluster 2 enriched DNA damage response and repair pathways (HMGB1, BIRC2, AKT3, PIK3R3, RICTOR) ^56–59^, consistent with a radioresistant, stem-like phenotype, while Cluster 3 showed suppression of apoptosis and immune signaling with oncogenic activation (AKT3, MAP3K1, PTPN11, RHOG, PLCG1) ^60–63^. Together, these findings show that high-dose irradiation (6 Gy) promotes a radioresistant, stem-like phenotype by suppressing apoptotic and immune signaling while activating DNA repair, chromatin remodeling, and oncogenic pathways, in contrast to standard fractionation (2 Gy), which primarily induces DNA repair and immune modulation.

Chromatin accessibility analysis revealed dose- and cluster-specific transcriptional suppression in glioblastoma stem-like cells following radiation. At 2 Gy, widespread closure occurred at TF binding sites linked to p53 signaling, including loss of over 1500 BATF sites, potentially relieving repression and enabling DNA damage responses. This pattern reversed at 6 Gy, where BATF binding increased in Clusters 0 and 1, correlating with p53 pathway suppression. Cluster-specific chromatin closure also affected immune and stress-related TFs: Clusters 0, 1, and 3 showed reduced accessibility at motifs regulating viral defense and biotic stress, while Cluster 2 showed suppression of interferon and oxidative stress pathways at 6 Gy. Notably, NFE2L2 and BATF emerged as key regulators across open and closed regions, suggesting their roles in modulating stress and immune response after radiation. These findings highlight distinct chromatin adaptations employed by each cluster in response to radiation dose.

In summary, our integrated single-cell transcriptomic and epigenetic analysis reveals that glioblastoma stem-like cells exhibit highly heterogeneous, dose- and cluster-specific responses to radiation. Each cluster engages distinct biological programs and regulatory mechanisms, shaped by gene-expression alterations and chromatin accessibility. By identifying dose- and cluster-specific vulnerabilities, our study lays the groundwork for precision radiotherapy strategies that target resistant subpopulations and exploit differential transcriptional states to improve glioblastoma treatment outcomes.

## RESOURCE AVAILABILITY

### Lead contact

Further information and requests for resources should be directed to and will be fulfilled by the lead contact, Aram Modrek.

### Materials availability

This study did not generate new reagents.

### Data and code availability

Single-cell RNA-seq and single-cell ATAC-seq data have been deposited at GEO as GSE309579.

## ACKNOWLEDGEMENTS

We are grateful for the support of all the funding agencies that made this work possible. Their contributions were essential to the completion of this study. This research is supported by National Institutes of Health – NCI grant K08CA263302, V Foundation Scholar Grant V2024-034, Donald E. and Delia B. Baxter Foundation Fellowship Award, and Achievement Research for College Scientists Foundation, Inc.- John H. Richardson Endowed Postdoctoral Fellowship Award. We also acknowledge the help from USC Norris Comprehensive Cancer Center- Molecular Genomics Core and its team, and the 10x Genomics team for their support during the study.

## AUTHOR CONTRIBUTIONS

Khoi Huynh designed the experiment with input from Aram Modrek. Khoi Huynh also performed all bioinformatics processing and analysis for the experiments. Sara Barcik Weissman performed all cell culture and 10x multiomic procedures with input and guidance from Aram Modrek and Cheol Park. USC genomic core was responsible for all sequencing of data. Blake Sun Change extracted the main biological functions for all pathways identified in the paper and generated heatmaps for the paper. Maria Quiloan contributed to manuscript refinement by reviewing grammar and layman’s logical structure during a revision round. David Tran, Eric Chang, and Kelly Street provided expert quality control on results and methods at appropriate steps.

## DECLARATION OF INTERESTS

The authors declare no competing interests.

### Declaration of generative AI and AI-assisted technologies in the writing process statement

During the preparation of this work, the author(s) used Microsoft Copilot to improve the readability and language of the manuscript. After using this tool/service, the author(s) reviewed and edited the content as needed and take(s) full responsibility for the content of the published article.

## SUPPLEMENTAL INFORMATION

**Figure S1.**
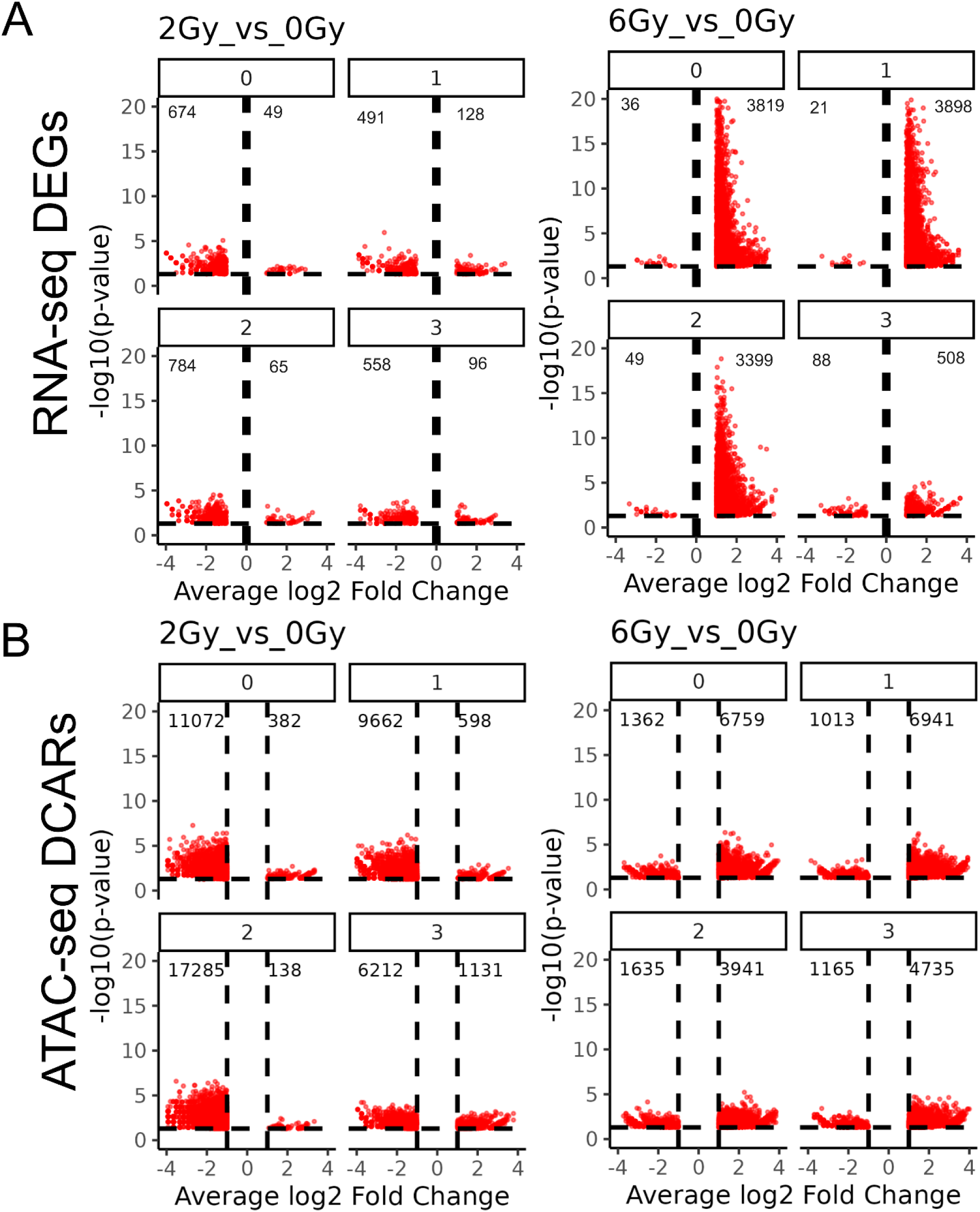
Volcano plots for DEGs. (A) Volcano plots showing differentially expressed genes (DEGs) for 2 Gy vs. 0 Gy (left) and 6 Gy vs. 0 Gy (right). Each point represents a gene, with log₂ fold change on the x-axis and –log₁₀ p-value on the y-axis. Genes with |log₂ fold change| ≥ 0.1 and p-value < 0.05 are highlighted as significantly differentially expressed (B) Volcano plots showing differentially accessible chromatin regions (DCARs) for 2 Gy vs. 0 Gy (left) and 6 Gy vs. 0 Gy (right). Each point represents a genomic region, with log₂ fold change in accessibility on the x-axis and –log₁₀ p-value on the y-axis. Regions with |log₂ fold change| ≥ 1 and p-value < 0.05 are highlighted as significantly differentially accessible.

**Figure S2.**
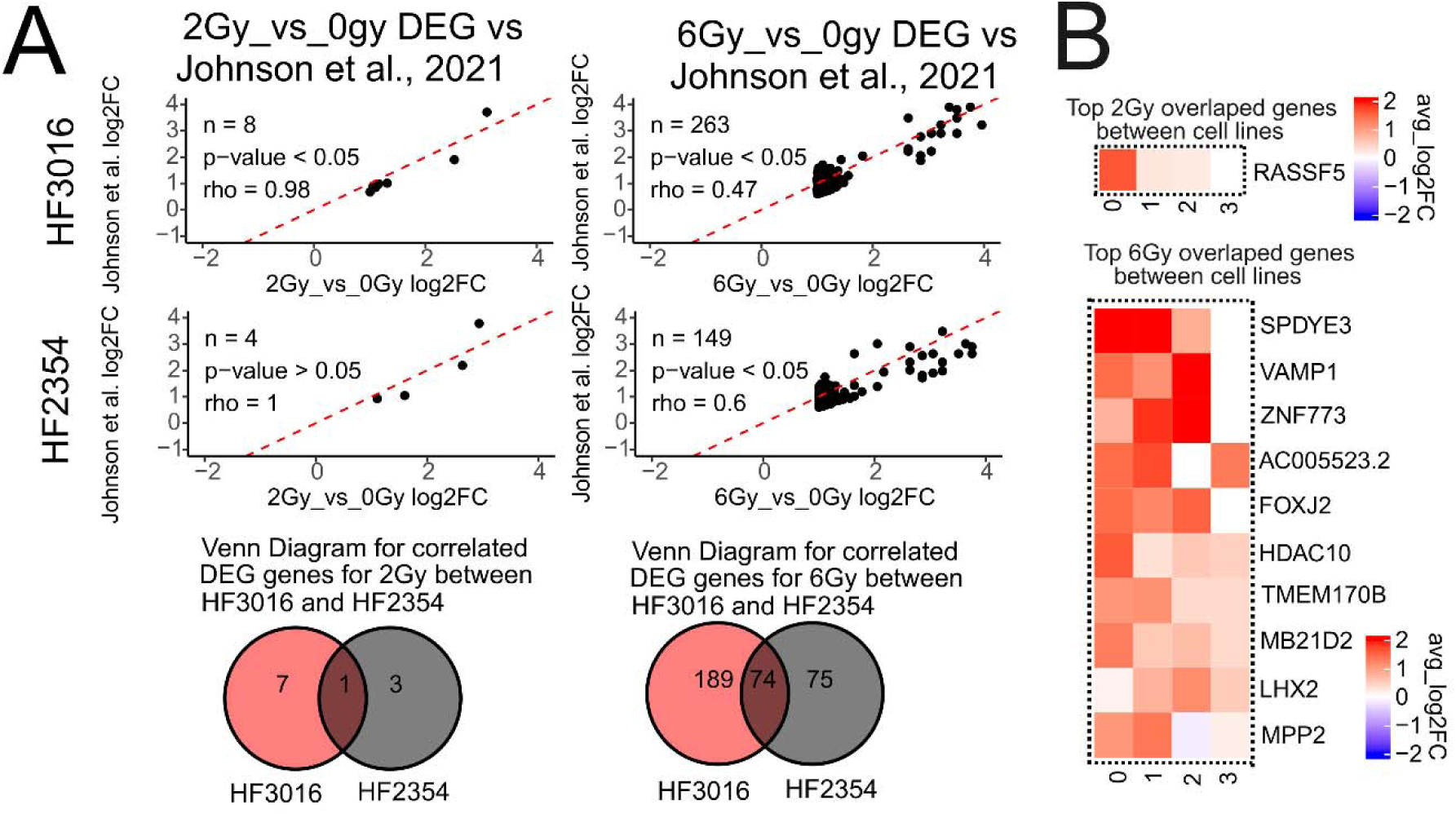
Conserved Radiation-Induced DEGs Across Cell Lines Independent of RNA Cluster Identity. (A) Volcano plots showing correlated differentially expressed genes (DEGs) between GSC20 and Johnson et al., 2021 HF2354 and HF3016 cell lines. Differential expression threshold using DESeq2 is p-value < 0.05. Correlation was assessed using Pearson correlation (p-value < 0.05) combined with differential expression criteria (p-value < 0.05). Bottom Venn diagrams illustrate the overlap of correlated DEGs in response to 2 Gy (left) and 6 Gy (right). (B) Heatmaps showing log2FoldChange values for the top five highest and lowest log2FoldChange of overlapping genes identified in panel A in response to 2 Gy (top) and to 6 Gy (bottom). Genes and samples are clustered using, and statistical significance for differential expressions were determined using DESeq2 with p-value < 0.05.

**Figure S3.**
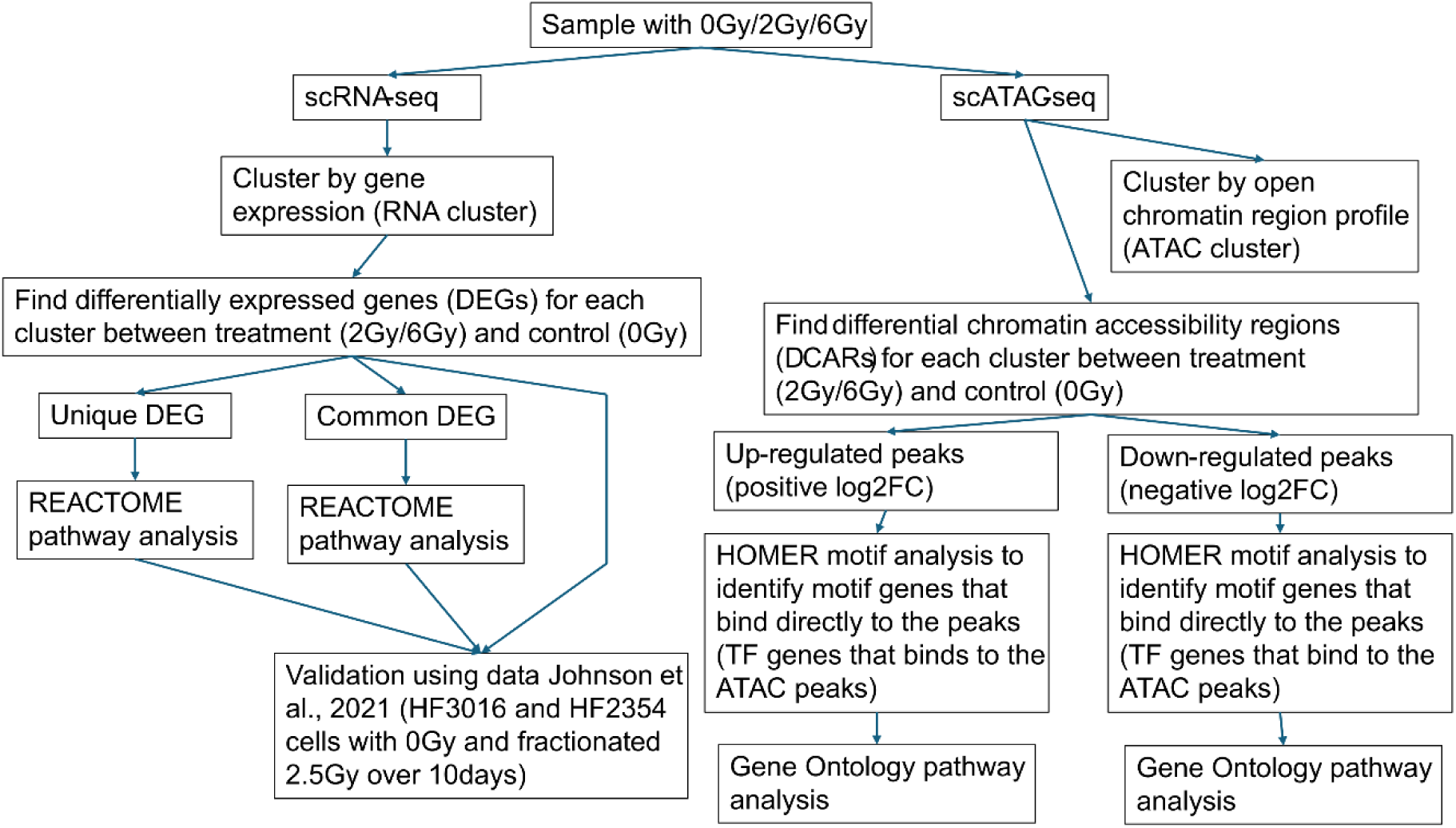
Experimental workflow.

**Document S1. Figure S1-S3 and Table S1-S4**

**Data S1. Differential gene expression across cell clusters and radiation dose conditions.**

**Data S2. Chromatin accessibility variation test across cell clusters and radiation exposure levels**

## STAR⍰METHODS

STAR<sup>®</sup> METHODS
| REAGENT OR RESOURCE | SOURCE | IDENTIFIER |
| --- | --- | --- |
| **Chemicals, peptides, and recombinant proteins** |  |  |
| Gibco B-27 supplement (50X, 10 mL) | Thermo Fisher Scientific | 17504044 |
| StemCell Technologies Human Recombinant bFGF (1,000 ug, ACF) | Stem Cell Technologies | 78134.2 |
| StemCell Technologies Human Recombinant EGF (1,000 ug, ACF) | Stem Cell Technologies | 78136.2 |
| Corning® DMEM/Hams F-12 50:50 Mix [+] L-glutamine (500 ml, 6/PK, Cat. No. 10-090-CV) | VWR International | 45000-344 |
| TRYPsin EDTA **0.25%** ST W/O CA/MAG (100mL, 6/CS, Cat. No. 25-053-CI) | VWR International | SC-45000-664 |
| **Critical commercial assays** |  |  |
| Chromium Next GEM Single Cell Multiome ATAC + Gene Expression Reagent Bundle, 16 rxns | 10x Genomics | PN-1000283 |
| **Deposited data** |  |  |
| scRNA-seq and scATAC-seq data | GEO | GSE309579 |
| **Experimental models: Cell lines** |  |  |
| GSC20 | Sulman Lab, New York University | N/a |
| **Software and algorithms** |  |  |
| AnnotationHub v3.14.0 | Morgan M et al. <sup>64</sup> | N/a |
| R v4.3.2 | R core team <sup>65</sup> | N/a |
| cellranger-arc v2.0.2 | 10x Genomics <sup>66</sup> | N/a |
| HOMER v5.1 | Heinz S et al. <sup>67</sup> | N/a |
| bedtools2 v2.30.0 | Quinlan AR et al. <sup>68</sup> | N/a |
| Samtools v1.17 | Heng Li et al. <sup>69</sup> | N/a |
| BSgenome.Hsapiens.UCSC.hg38 v1.4.5 | Team TBD <sup>70</sup> | N/a |
| EnhancedVolcano v1.24.0 | Blighe K et al. <sup>71</sup> | N/a |
| EnsDb.Hsapiens.v86 v2.99.0 | Rainer J <sup>72</sup> | N/a |
| JASPAR2020 v0.99.10 | Fornes et al. <sup>73</sup> | N/a |
| ReactomePA v1.50.0 | GuangChuang Y et al. <sup>74</sup> | N/a |
| Seurat v5.2.0 | Hao Y et al. <sup>11</sup> | N/a |
| SeuratDisk v0.0.0.9021 | Hoffman P, SeuratDisk, https://github.com/mojaveazure/seurat-disk | N/a |
| Signac v1.14.0 | Stuart T et al. <sup>75</sup> | N/a |
| TFBSTools v1.44.0 | Tan G et al. <sup>76</sup> | N/a |
| biomaRt v2.62.1 | Durinck S et al. <sup>77</sup> | N/a |
| Cowplot v1.1.3 | Wilke C, Cowplot, https://wilkelab.org/cowplot/reference/cowplot.html | N/a |
| Fgsea v1.32.4 | Korotkevich G et al. <sup>78</sup> | N/a |
| ggplot2 3.5.2 | Wickham H et al. <sup>79</sup> | N/a |
| Ggpubr v0.6.0 | Kassambara A et al. <sup>80</sup> | N/a |
| Msigdbr v10.0.2 | Bhuva DA et al. <sup>81</sup> | N/a |
| org.Hs.eg.db v3.20.0 | Carlson M <sup>82</sup> | N/a |
| Pathwork v1.3.0.9000 | Pedersen T, pathwork, https://patchwork.data-imaginist.com/ | N/a |
| Tidyverse v2.0.0 | Wickham H et al. <sup>83</sup> | N/a |
| Scales v1.4.0 | Wickham H et al. <sup>84</sup> | N/a |

## METHOD DETAILS

A brief experimental workflow can be found in Figure S3.

### Glioma Stem-like Cell cultures and radiation treatment

GSC20 cells were from the Sulman Lab group and previously published in ^13^. All cells are cultured in standard neural basal medium (NBM), which consists of Corning DMEM/Hams F-12 50/50 mix, 1x B27 supplement, 20ng/mL EGF, 20ng/mL bFGF, and 1x anti-mycotic anti-bacterial solution. To irradiate the cell cultures, both the control and treatment cell culture flasks are brought to the irradiator. Irradiation is done using X-RAD 320 with kV at 250.0, mA at 16.0, doses at 200 (2 Gy) and at 600 (6 Gy), fill at 1, and SSD at 50cm.

### Single-nucleus multi-omics sequencing

Cells were harvested, and nuclei were isolated following the 10x Genomics protocol. Nuclei were processed using the Chromium Single Cell Multiome ATAC + Gene Expression kit and the Chromium Controller (10x Genomics), according to the manufacturer’s instructions. Approximately 5000-10000 nuclei were loaded per channel with the multiome gel bead. Library quantification was performed using a Qubit Fluorometer, and fragment profiles were assessed with a Fragment Analyzer. Sequencing of scRNA-seq and scATAC-seq libraries was carried out on a NextSeq2000 P2 platform using recommended read depths of 25000 read pairs per nucleus.

### scRNA-seq and scATAC-seq processing

Both scRNA-seq and scATAC-seq datasets were demultiplexed, processed, and aligned using Cell Ranger ARC with the human reference genome hg38. Following alignment, the datasets were analyzed in R using the Seurat package for scRNA-seq and the Signac package for scATAC-seq. Peak calling for the ATAC-seq data was performed using MACS3 on the hg38 genome, excluding regions listed in the genomic blacklist. RNA-seq complexity was calculated by dividing the number of RNA reads per gene by the total RNA read count. Cells were then filtered based on several quality control metrics: total RNA read counts were required to be between 500 and 50000, and total ATAC read counts between 500 and 100000. Additionally, RNA complexity had to exceed 0.8, nucleosome signal (calculated via Seurat’s NucleosomeSignal) had to be less than 1, and transcription start site (TSS) enrichment (calculated via Seurat’s TSSenrichment) had to be greater than 1. For RNA-seq data, normalization was done using Seurat SCTransform. Following normalization, RNA data were conFigured to use the "SCT" assay as the default, and variable features were set using all features from the SCT assay. Principal component analysis (PCA) was performed to reduce dimensionality, followed by Uniform Manifold Approximation and Projection (UMAP) using the first 20 principal components to visualize the data.

Subsequently, a nearest-neighbor graph was constructed based on these components, and clustering was performed at a resolution of 0.4 to identify distinct cell populations. For ATAC-seq data, term Frequency–Inverse Document Frequency (TF-IDF) normalization was applied, followed by selection of top features. Singular Value Decomposition (SVD) was then performed using 20 components, excluding the first dimension, which typically correlates with sequencing depth.

### scRNA-seq and scATAC-seq differential analysis

To perform differential gene expression analysis, the Seurat object was first switched to the "SCT" assay. For each RNA-defined cluster, differential gene expression analysis was conducted using Seurat’s FindMarkers function, applying a significance threshold of p < 0.05 and a log₂ fold change cutoff of ±0.1. In parallel, differential accessibility analysis was performed to identify polymorphic chromatin regions (ATAC-seq peaks) associated with each RNA cluster. This was done using the same FindMarkers function but applied to the "peaks" assay of the Seurat object with log₂ fold change cutoff of ±1 instead.

### scRNA-seq gene set enrichment analysis (GSEA)

Differentially expressed genes identified for each RNA cluster were subjected to Gene Set Enrichment Analysis (GSEA) to uncover biological pathways affected by radiation exposure. GSEA was performed using the fgsea package, leveraging gene-level log₂ fold change values and querying across all MSigDB databases via custom scripts. Enriched pathways were filtered to retain only those with a p-value less than 0.05. Additionally, each identified pathway was annotated with a corresponding top-level biological function using a custom script that mapped MSigDB terms to broader functional categories.

### scATAC-seq motif analysis and motif gene set (transcription factor gene enrichment analysis (GSEA)

Differentially accessible chromatin regions identified using Seurat’s FindMarkers function were categorized into "open" (log₂ fold change > 1) and "closed" (log₂ fold change < -1) peaks. Each set of peaks was submitted to HOMER for motif enrichment analysis using a fixed window size of 200 bp. From the HOMER output, only transcription factor (TF) motifs with a binding site located directly within the differentially accessible region (distance = 0) were retained. These motif-associated TFs were then used for gene set enrichment analysis (GSEA), following the same approach described previously. Instead of using log₂ fold change values, the number of binding sites per motif gene was used to assess biological functions affected by radiation within each RNA-defined cluster.

### Differentially expressed genes validation

To validate identified DEGs, we downloaded data from Johnson et al. 2021 ^85^. This is scRNA-seq on cells receiving 0 Gy (control), and fractionated 2.5 Gy four times (treatment). The RNA data sets are processed, normalized, and analyzed similarly to what we had done for our data sets. However, instead of performing clustering, we projected our RNA clusters onto Johnson data sets using gene expression profiles. Then, we compared our DEGs to their identified DEGs for every RNA cluster while keeping all DEGs within 60% of each other in terms of log_2_FC magnitude and to have the same direction of regulation (same sign of log_2_FC). Then, we performed Pearson correlation on those DEGs that appear in both our data sets and Johnson’s data sets. Furthermore, we also performed GSEA on correlated DEGs.

## REFERENCE

1. Stupp, R., Taillibert, S., Kanner, A., Read, W., Steinberg, D.M., Lhermitte, B., Toms, S., Idbaih, A., Ahluwalia, M.S., Fink, K., et al. (2017). Effect of tumor-treating fields plus maintenance temozolomide vs maintenance temozolomide alone on survival in patients with glioblastoma. JAMA 318, 2306.

2. Johnson, A.L., and Lopez-Bertoni, H. (2024). Cellular diversity through space and time: adding new dimensions to GBM therapeutic development. Front. Genet. 15, 1356611.

3. Crivii, C.-B., Bo□ca, A.B., Melincovici, C.S., Constantin, A.-M., Mărginean, M., Dronca, E., Sufle□el, R., Gonciar, D., Bungărdean, M., and □ovrea, A. (2022). Glioblastoma microenvironment and cellular interactions. Cancers (Basel) 14, 1092.

4. Wang, X., Sun, Q., Liu, T., Lu, H., Lin, X., Wang, W., Liu, Y., Huang, Y., Huang, G., Sun, H., et al. (2024). Single-cell multi-omics sequencing uncovers region-specific plasticity of glioblastoma for complementary therapeutic targeting. Sci. Adv. 10, eadn4306.

5. Eisenbarth, D., and Wang, Y.A. (2023). Glioblastoma heterogeneity at single cell resolution. Oncogene 42, 2155–2165.

6. Singh, S., Dey, D., Barik, D., Mohapatra, I., Kim, S., Sharma, M., Prasad, S., Wang, P., Singh, A., and Singh, G. (2025). Glioblastoma at the crossroads: current understanding and future therapeutic horizons. Signal Transduct. Target. Ther. 10, 213.

7. Rabah, N., Ait Mohand, F.-E., and Kravchenko-Balasha, N. (2023). Understanding glioblastoma signaling, heterogeneity, invasiveness, and drug delivery barriers. Int. J. Mol. Sci. 24. 10.3390/ijms241814256.

8. Nomura, M., Spitzer, A., Johnson, K.C., Garofano, L., Nehar-Belaid, D., Galili Darnell, N., Greenwald, A.C., Bussema, L., Oh, Y.T., Varn, F.S., et al. (2025). The multilayered transcriptional architecture of glioblastoma ecosystems. Nat. Genet. 57, 1155–1167.

9. Kondo, T. (2022). Glioblastoma-initiating cell heterogeneity generated by the cell-of-origin, genetic/epigenetic mutation and microenvironment. Semin. Cancer Biol. 82, 176–183.

10. Billoir, M., Crepin, D., Plaszczynski, S., Grammaticos, B., Seksek, O., and Badoual, M. (2025). The temporal response of a glioma cell population to irradiation: modelling the effect of dose and cell density. R. Soc. Open Sci. 12, 241917.

11. Hao, Y., Stuart, T., Kowalski, M.H., Choudhary, S., Hoffman, P., Hartman, A., Srivastava, A., Molla, G., Madad, S., Fernandez-Granda, C., et al. (2024). Dictionary learning for integrative, multimodal and scalable single-cell analysis. Nat. Biotechnol. 42, 293–304.

12. Verhaak, R.G.W., Hoadley, K.A., Purdom, E., Wang, V., Qi, Y., Wilkerson, M.D., Miller, C.R., Ding, L., Golub, T., Mesirov, J.P., et al. (2010). Integrated genomic analysis identifies clinically relevant subtypes of glioblastoma characterized by abnormalities in PDGFRA, IDH1, EGFR, and NF1. Cancer Cell 17, 98–110.

13. Bhat, K.P.L., Balasubramaniyan, V., Vaillant, B., Ezhilarasan, R., Hummelink, K., Hollingsworth, F., Wani, K., Heathcock, L., James, J.D., Goodman, L.D., et al. (2013). Mesenchymal differentiation mediated by NF-κB promotes radiation resistance in glioblastoma. Cancer Cell 24, 331–346.

14. Moreno, M., Pedrosa, L., Paré, L., Pineda, E., Bejarano, L., Martínez, J., Balasubramaniyan, V., Ezhilarasan, R., Kallarackal, N., Kim, S.-H., et al. (2017). GPR56/ADGRG1 inhibits mesenchymal differentiation and radioresistance in glioblastoma. Cell Rep. 21, 2183–2197.

15. Behnan, J., Finocchiaro, G., and Hanna, G. (2019). The landscape of the mesenchymal signature in brain tumours. Brain 142, 847–866.

16. Fedele, M., Cerchia, L., Pegoraro, S., Sgarra, R., and Manfioletti, G. (2019). Proneural-mesenchymal transition: Phenotypic plasticity to acquire multitherapy resistance in glioblastoma. Int. J. Mol. Sci. 20, 2746.

17. Kim, Y., Varn, F.S., Park, S.-H., Yoon, B.W., Park, H.R., Lee, C., Verhaak, R.G.W., and Paek, S.H. (2021). Perspective of mesenchymal transformation in glioblastoma. Acta Neuropathol. Commun. 9, 50.

18. Mahdi, A., Aittaleb, M., and Tissir, F. (2025). Targeting glioma stem cells: Therapeutic opportunities and challenges. Cells 14, 675.

19. Spehalski, E.I., Lee, J.A., Peters, C., Tofilon, P., and Camphausen, K. (2019). The quiescent metabolic phenotype of glioma stem cells. J. Proteomics Bioinform. 12, 96–103.

20. Visioli, A., Trivieri, N., Mencarelli, G., Giani, F., Copetti, M., Palumbo, O., Pracella, R., Cariglia, M.G., Barile, C., Mischitelli, L., et al. (2023). Different states of stemness of glioblastoma stem cells sustain glioblastoma subtypes indicating novel clinical biomarkers and high-efficacy customized therapies. J. Exp. Clin. Cancer Res. 42, 244.

21. Babačić, H., Galardi, S., Umer, H.M., Hellström, M., Uhrbom, L., Maturi, N., Cardinali, D., Pellegatta, S., Michienzi, A., Trevisi, G., et al. (2023). Glioblastoma stem cells express non-canonical proteins and exclusive mesenchymal-like or non-mesenchymal-like protein signatures. Mol. Oncol. 17, 238–260.

22. Lo Cascio, C., Luna Melendez, E., McNamara, J., Schultz, R., Tien, A.-C., Plaisier, C., and Mehta, S. (2019). Stem-15. Radiation-induced modulation of neurodevelopmental pathways in Classical glioma stem cells. Neuro. Oncol. 21, vi236–vi237.

23. Skouras, P., Markouli, M., Papadatou, I., and Piperi, C. (2024). Targeting epigenetic mechanisms of resistance to chemotherapy in gliomas. Crit. Rev. Oncol. Hematol. 204, 104532.

24. Wang, Z., Zhang, H., Xu, S., Liu, Z., and Cheng, Q. (2021). The adaptive transition of glioblastoma stem cells and its implications on treatments. Signal Transduct. Target. Ther. 6, 124.

25. Tian, G., Song, Y., Zhang, Y., Kan, L., Hou, A., and Han, S. (2025). Phenotypic variations in glioma stem cells: regulatory mechanisms and implications for therapeutic strategies. J. Transl. Med. 23, 984.

26. Brackmann, L.K., Poplawski, A., Grandt, C.L., Schwarz, H., Hankeln, T., Rapp, S., Zahnreich, S., Galetzka, D., Schmitt, I., Grad, C., et al. (2020). Comparison of time and dose dependent gene expression and affected pathways in primary human fibroblasts after exposure to ionizing radiation. Mol. Med. 26. 10.1186/s10020-020-00203-0.

27. Wang, Z., Zang, C., Cui, K., Schones, D.E., Barski, A., Peng, W., and Zhao, K. (2009). Genome-wide mapping of HATs and HDACs reveals distinct functions in active and inactive genes. Cell 138, 1019–1031.

28. Gong, F., and Miller, K.M. (2013). Mammalian DNA repair: HATs and HDACs make their mark through histone acetylation. Mutat. Res. 750, 23–30.

29. Wang, X., and Zhao, J. (2022). Targeted cancer therapy based on acetylation and deacetylation of key proteins involved in double-strand break repair. Cancer Manag. Res. 14, 259–271.

30. Verza, F.A., Das, U., Fachin, A.L., Dimmock, J.R., and Marins, M. (2020). Roles of histone deacetylases and inhibitors in anticancer therapy. Cancers (Basel) 12, 1664.

31. Xu, S., Höglund, M., Hâkansson, L., and Venge, P. (2000). Granulocyte colony-stimulating factor (G-CSF) induces the production of cytokines in vivo. Br. J. Haematol. 108, 848–853.

32. Zhang, Q., Jiang, L., Wang, W., Huber, A.K., Valvo, V.M., Jungles, K.M., Holcomb, E.A., Pearson, A.N., The, S., Wang, Z., et al. (2024). Potentiating the radiation-induced type I interferon antitumoral immune response by ATM inhibition in pancreatic cancer. JCI Insight 9. 10.1172/jci.insight.168824.

33. Goedegebuure, R.S.A., Kleibeuker, E.A., Buffa, F.M., Castricum, K.C.M., Haider, S., Schulkens, I.A., Ten Kroode, L., van den Berg, J., Jacobs, M.A.J.M., van Berkel, A.-M., et al. (2021). Interferon- and STING-independent induction of type I interferon stimulated genes during fractionated irradiation. J. Exp. Clin. Cancer Res. 40, 161.

34. Ma, W., Huang, G., Wang, Z., Wang, L., and Gao, Q. (2023). IRF7: role and regulation in immunity and autoimmunity. Front. Immunol. 14, 1236923.

35. Rübe, C.E., Al-Razaq, M.A.A., Meier, C., Hecht, M., and Rübe, C. (2025). Role of ionizing radiation in shaping the complex multi-layered epigenome. Epigenomes 9, 29.

36. Gupta, A., Hunt, C.R., Chakraborty, S., Pandita, R.K., Yordy, J., Ramnarain, D.B., Horikoshi, N., and Pandita, T.K. (2014). Role of 53BP1 in the regulation of DNA double-strand break repair pathway choice. Radiat. Res. 181, 1–8.

37. Sokolov, M.V., Smilenov, L.B., Hall, E.J., Panyutin, I.G., Bonner, W.M., and Sedelnikova, O.A. (2005). Ionizing radiation induces DNA double-strand breaks in bystander primary human fibroblasts. Oncogene 24, 7257–7265.

38. Hill, M.A., and Ullrich, R.L. (2019). Ionizing radiation. In Tumour Site Concordance and Mechanisms of Carcinogenesis (International Agency for Research on Cancer).

39. Dahl, H., Ballangby, J., Tengs, T., Wojewodzic, M.W., Eide, D.M., Brede, D.A., Graupner, A., Duale, N., and Olsen, A.-K. (2023). Dose rate dependent reduction in chromatin accessibility at transcriptional start sites long time after exposure to gamma radiation. Epigenetics 18, 2193936.

40. Peng, Q., Weng, K., Li, S., Xu, R., Wang, Y., and Wu, Y. (2021). A perspective of epigenetic regulation in radiotherapy. Front. Cell Dev. Biol. 9, 624312.

41. Kumar, R., Horikoshi, N., Singh, M., Gupta, A., Misra, H.S., Albuquerque, K., Hunt, C.R., and Pandita, T.K. (2012). Chromatin modifications and the DNA damage response to ionizing radiation. Front. Oncol. 2, 214.

42. Johnson, K.C., Anderson, K.J., Courtois, E.T., Gujar, A.D., Barthel, F.P., Varn, F.S., Luo, D., Seignon, M., Yi, E., Kim, H., et al. (2021). Single-cell multimodal glioma analyses identify epigenetic regulators of cellular plasticity and environmental stress response. Nat. Genet. 53, 1456–1468.

43. Xue, S., Luo, Z., Mao, Y., and Liu, S. (2025). A comprehensive analysis of the pyruvate kinase M1/2 (PKM) in human cancer. Gene 937, 149155.

44. Alexandrou, A.T., Duan, Y., Xu, S., Tepper, C., Fan, M., Tang, J., Berg, J., Basheer, W., Valicenti, T., Wilson, P.F., et al. (2022). PERIOD 2 regulates low-dose radioprotection via PER2/pGSK3β/β-catenin/Per2 loop. iScience 25, 105546.

45. Zhang, P., Chen, X., Zhang, L., Cao, D., Chen, Y., Guo, Z., and Chen, J. (2022). POLE2 facilitates the malignant phenotypes of glioblastoma through promoting AURKA-mediated stabilization of FOXM1. Cell Death Dis. 13, 61.

46. Richardson, T.E., Sathe, A.A., Kanchwala, M., Jia, G., Habib, A.A., Xiao, G., Snuderl, M., Xing, C., and Hatanpaa, K.J. (2018). Genetic and epigenetic features of rapidly progressing IDH-mutant astrocytomas. J. Neuropathol. Exp. Neurol. 77, 542–548.

47. Feng, T., Zheng, L., Liu, F., Xu, X., Mao, S., Wang, X., Liu, J., Lu, Y., Zhao, W., Yu, X., et al. (2016). Growth factor progranulin promotes tumorigenesis of cervical cancer via PI3K/Akt/mTOR signaling pathway. Oncotarget 7, 58381–58395.

48. Lau, M.-T., Klausen, C., and Leung, P.C.K. (2011). E-cadherin inhibits tumor cell growth by suppressing PI3K/Akt signaling via β-catenin-Egr1-mediated PTEN expression. Oncogene 30, 2753–2766.

49. Wang, H., Guan, Q., Nan, Y., Ma, Q., and Zhong, Y. (2019). Overexpression of human MX2 gene suppresses cell proliferation, migration, and invasion via ERK/P38/NF-κB pathway in glioblastoma cells. J. Cell. Biochem. 120, 18762–18770.

50. Li, Z., Qian, R., Li, M., Li, J., Guo, Y., Zhou, Y., and Ma, C. (2025). HERC5/ISG15 Enhances Glioblastoma Stemness and Tumor Progression by mediating SERBP1protein stability. Neuromolecular Med. 27, 7.

51. Blomquist, M.R., Eghlimi, R., Beniwal, A., Grief, D., Nascari, D.G., Inge, L., Sereduk, C.P., Tuncali, S., Roos, A., Inforzato, H., et al. (2024). EGFRvIII confers sensitivity to saracatinib in a STAT5-dependent manner in glioblastoma. Int. J. Mol. Sci. 25, 6279.

52. Zhao, J., Zhang, L., Dong, X., Liu, L., Huo, L., and Chen, H. (2018). High expression of vimentin is associated with progression and a poor outcome in glioblastoma. Appl. Immunohistochem. Mol. Morphol. 26, 337–344.

53. Xu, J., Zhang, J., Chen, W., and Ni, X. (2024). The tumor-associated fibrotic reactions in microenvironment aggravate glioma chemoresistance. Front. Oncol. 14, 1388700.

54. Westhoff, M.-A., Kandenwein, J.A., Karl, S., Vellanki, S.H.K., Braun, V., Eramo, A., Antoniadis, G., Debatin, K.-M., and Fulda, S. (2009). The pyridinylfuranopyrimidine inhibitor, PI-103, chemosensitizes glioblastoma cells for apoptosis by inhibiting DNA repair. Oncogene 28, 3586–3596.

55. Zhu, R., Ye, X., Lu, X., Xiao, L., Yuan, M., Zhao, H., Guo, D., Meng, Y., Han, H., Luo, S., et al. (2025). ACSS2 acts as a lactyl-CoA synthetase and couples KAT2A to function as a lactyltransferase for histone lactylation and tumor immune evasion. Cell Metab. 37, 361–376.e7.

56. Qiu, H., Shao, Z., Wen, X., Qu, D., Liu, Z., Chen, Z., Zhang, X., Ding, X., and Zhang, L. (2024). HMGB1/TREM2 positive feedback loop drives the development of radioresistance and immune escape of glioblastoma by regulating TLR4/Akt signaling. J. Transl. Med. 22, 688.

57. Vidomanova, E., Majercikova, Z., Dibdiakova, K., Pilchova, I., Racay, P., and Hatok, J. (2022). The effect of temozolomide on apoptosis-related gene expression changes in glioblastoma cells. Bratisl. Lek. Listy 123, 236–243.

58. Wang, S., Ming, H., Wang, Z., Zhai, X., Zhang, X., Wu, D., Bo, Y., Wang, H., Luo, Y., Han, Z., et al. (2025). NONHSAT141192.2 facilitates the stemness and radioresistance of glioma stem cells via the regulation of PIK3R3 and SOX2. CNS Neurosci. Ther. 31, e70279.

59. Ramaiah, M.J., and Kumar, K.R. (2021). mTOR-Rictor-EGFR axis in oncogenesis and diagnosis of glioblastoma multiforme. Mol. Biol. Rep. 48, 4813–4835.

60. Wang, L.-B., Karpova, A., Gritsenko, M.A., Kyle, J.E., Cao, S., Li, Y., Rykunov, D., Colaprico, A., Rothstein, J.H., Hong, R., et al. (2021). Proteogenomic and metabolomic characterization of human glioblastoma. Cancer Cell 39, 509–528.e20.

61. Kwiatkowska, A., Didier, S., Fortin, S., Chuang, Y., White, T., Berens, M.E., Rushing, E., Eschbacher, J., Tran, N.L., Chan, A., et al. (2012). The small GTPase RhoG mediates glioblastoma cell invasion. Mol. Cancer 11, 65.

62. Zhou, S., Niu, R., Sun, H., Kim, S.-H., Jin, X., and Yin, J. (2022). The MAP3K1/c-JUN signaling axis regulates glioblastoma stem cell invasion and tumor progression. Biochem. Biophys. Res. Commun. 612, 188–195.

63. Paul-Samojedny, M., Suchanek, R., Borkowska, P., Pudełko, A., Owczarek, A., Kowalczyk, M., Machnik, G., Fila-Daniłow, A., and Kowalski, J. (2014). Knockdown of AKT3 (PKBγ) and PI3KCA suppresses cell viability and proliferation and induces the apoptosis of glioblastoma multiforme T98G cells. Biomed Res. Int. 2014, 768181.

64. Martin Morgan [cre], Marc Carlson [ctb], Dan Tenenbaum [ctb], Sonali Arora [ctb] (2017). AnnotationHub (Bioconductor) 10.18129/B9.BIOC.ANNOTATIONHUB.

65. Computing, R., and Others (2013). R: A language and environment for statistical computing. R Core Team.

66. Zheng, G.X.Y., Terry, J.M., Belgrader, P., Ryvkin, P., Bent, Z.W., Wilson, R., Ziraldo, S.B., Wheeler, T.D., McDermott, G.P., Zhu, J., et al. (2017). Massively parallel digital transcriptional profiling of single cells. Nat. Commun. 8, 14049.

67. Heinz, S., Benner, C., Spann, N., Bertolino, E., Lin, Y.C., Laslo, P., Cheng, J.X., Murre, C., Singh, H., and Glass, C.K. (2010). Simple combinations of lineage-determining transcription factors prime cis-regulatory elements required for macrophage and B cell identities. Mol. Cell 38, 576–589.

68. Quinlan, A.R., and Hall, I.M. (2010). BEDTools: a flexible suite of utilities for comparing genomic features. Bioinformatics 26, 841–842.

69. 69. Li, H., Bob, H., Wysoker, A., Fennell, T., Ruan, J., Homer, N., Marth, G., Abecasis, G., and Durbin, R. (2009). The Sequence Alignment/Map (SAM) Format and.

70. The Bioconductor Dev Team (2017). BSgenome.Hsapiens.UCSC.hg38 (Bioconductor) 10.18129/B9.BIOC.BSGENOME.HSAPIENS.UCSC.HG38.

71. Blighe, K. (2018). EnhancedVolcano (Bioconductor) 10.18129/B9.BIOC.ENHANCEDVOLCANO.

72. Rainer, J. (2017). EnsDb.Hsapiens.v86 (Bioconductor) 10.18129/B9.BIOC.ENSDB.HSAPIENS.V86.

73. Fornes, O., Castro-Mondragon, J.A., Khan, A., van der Lee, R., Zhang, X., Richmond, P.A., Modi, B.P., Correard, S., Gheorghe, M., Baranašić, D., et al. (2020). JASPAR 2020: update of the open-access database of transcription factor binding profiles. Nucleic Acids Res. 48, D87–D92.

74. Guangchuang Yu [aut, cre], Vladislav Petyuk [ctb] (2017). ReactomePA (Bioconductor) 10.18129/B9.BIOC.REACTOMEPA.

75. Stuart, T., Srivastava, A., Madad, S., Lareau, C.A., and Satija, R. (2021). Single-cell chromatin state analysis with Signac. Nat. Methods 18, 1333–1341.

76. Ge Tan [aut, C. (2017). TFBSTools (Bioconductor) 10.18129/B9.BIOC.TFBSTOOLS.

77. Steffen Durinck, Wolfgang Huber (2017). biomaRt (Bioconductor) 10.18129/B9.BIOC.BIOMART.

78. Alexey Sergushichev [aut, C. (2017). fgsea (Bioconductor) 10.18129/B9.BIOC.FGSEA.

79. 79. Wickham, H., Chang, W., Henry, L., Pedersen, T.L., Takahashi, K., Wilke, C., Woo, K., Yutani, H., Dunnington, D., and van den Brand, T. (2007). Ggplot2: Create elegant data visualisations using the grammar of graphics. (The R Foundation). 10.32614/cran.package.ggplot2 https://doi.org/10.32614/cran.package.ggplot2.

80. Kassambara, A. (2016). ggpubr: “ggplot2” Based Publication Ready Plots. (The R Foundation). 10.32614/cran.package.ggpubr https://doi.org/10.32614/cran.package.ggpubr.

81. Dharmesh D. Bhuva [aut, cre] (<https://orcid.org/0000-0002-6398-9157>), Gordon K. Smyth [aut] (<https://orcid.org/0000-0001-9221-2892>), Alexandra Garnham [aut] (<https://orcid.org/0000-0002-8312-8450>) (2021). msigdb (Bioconductor) 10.18129/B9.BIOC.MSIGDB.

82. Carlson, M. (2017). org.Hs.eg.db (Bioconductor) 10.18129/B9.BIOC.ORG.HS.EG.DB.

83. Wickham, H., Averick, M., Bryan, J., Chang, W., McGowan, L., François, R., Grolemund, G., Hayes, A., Henry, L., Hester, J., et al. (2019). Welcome to the tidyverse. J. Open Source Softw. 4, 1686.

84. Wickham, H., Pedersen, T.L., and Seidel, D. (2011). Scales: Scale functions for visualization. (The R Foundation). 10.32614/cran.package.scales https://doi.org/10.32614/cran.package.scales.

85. Johnson, K.C., Anderson, K.J., Courtois, E.T., Barthel, F.P., Varn, F.S., Luo, D., Seignon, M., Yi, E., Kim, H., Estecio, M.R.H., et al. (2020). Single-cell multimodal glioma analyses reveal epigenetic regulators of cellular plasticity and environmental stress response. bioRxiv. 10.1101/2020.07.22.215335.

