## Supplementary material for "Single-Cell Multiomic Profiling Uncovers Radiation Dosage-Sensitive, Cluster-Specific Regulatory Dynamics in Glioblastoma": Document S1

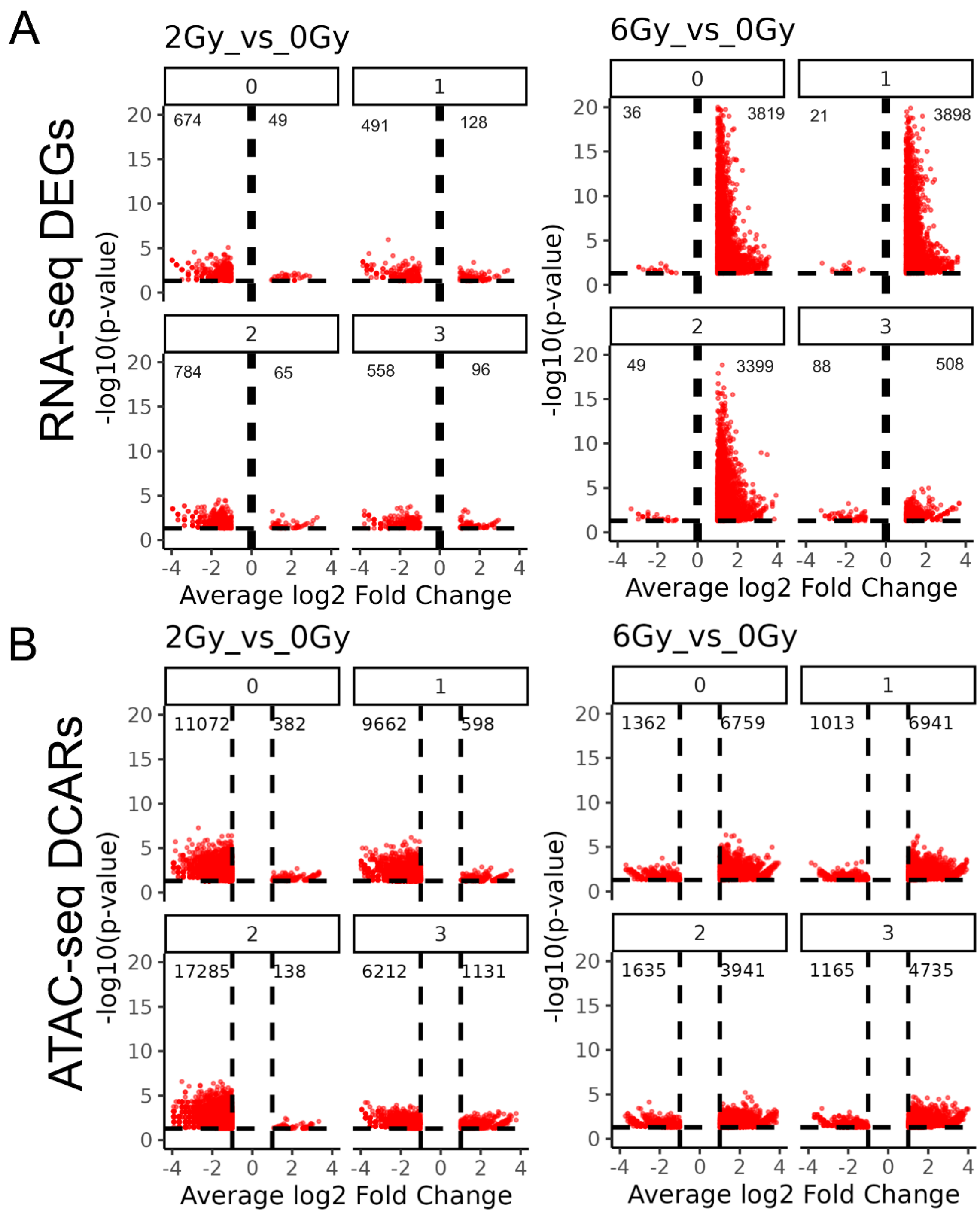


**Figure S1 Volcano plots for DEGs**

(A) Volcano plots showing differentially expressed genes (DEGs) for 2 Gy vs. 0 Gy (left) and 6 Gy vs. 0 Gy (right). Each point represents a gene, with $\log_2$ fold change on the x-axis and $-\log_{10}$ p-value on the y-axis. Genes with $|\log_2$ fold change$| \geq 0.1$ and p-value $< 0.05$ are highlighted as significantly differentially expressed (B) Volcano plots showing differentially accessible chromatin regions (DACRs) for 2 Gy vs. 0 Gy (left) and 6 Gy vs. 0 Gy (right). Each point represents a genomic region, with $\log_2$ fold change in accessibility on the x-axis and $-\log_{10}$ p-value on the y-axis. Regions with $|\log_2$ fold change$| \geq$ and p-value $< 0.05$ are highlighted as significantly differentially accessible.

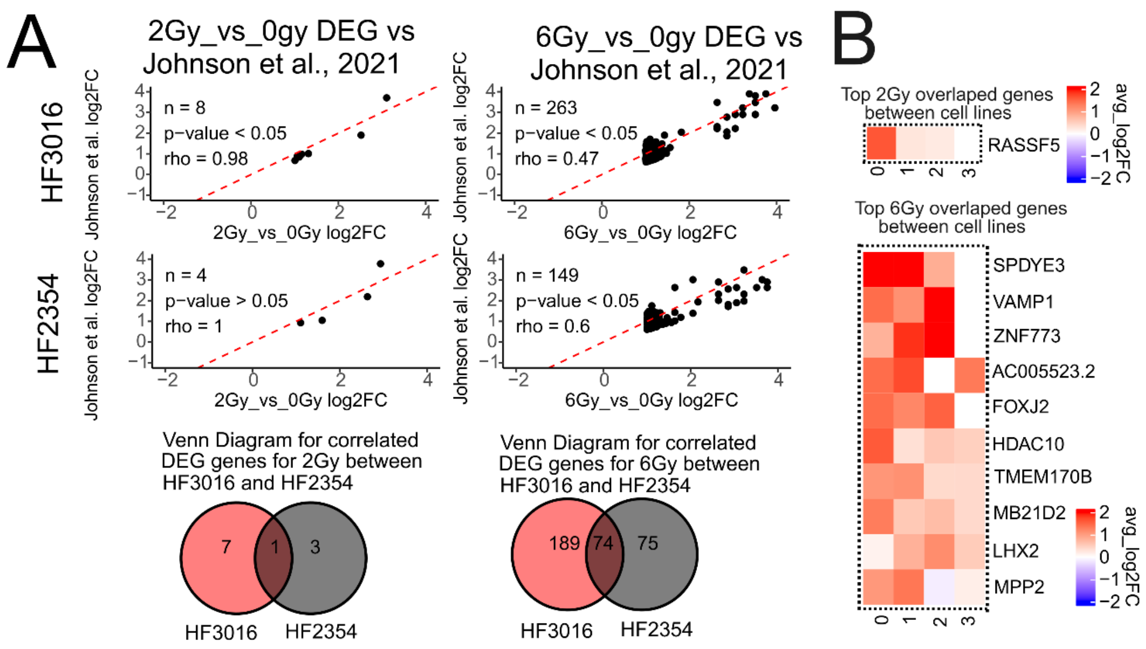


**Figure S2 Conserved Radiation-Induced DEGs Across Cell Lines Independent of RNA Cluster Identity**

(A) Volcano plots showing correlated differentially expressed genes (DEGs) between GSC20 and Johnson et al., 2021 HF2354 and HF3016 cell lines. Each point represents a gene, with $\log_2$ fold change on the x-axis and $-\log_{10}$ p-value on the y-axis. Genes with $|\log_2$ fold change$| \geq 0.1$ and p-value < 0.05 are highlighted as significantly correlated if they have the same $\log_2$FC sign. Venn diagrams illustrate the overlap of correlated DEGs in response to 2 Gy (left) and 6 Gy (right). (B) Heatmaps showing normalized expression (z-score) of overlapping genes identified in panel A across all conditions. Genes and samples are clustered using Euclidean distance, and statistical significance for differential expression was determined using DESeq2 with p-value < 0.05.

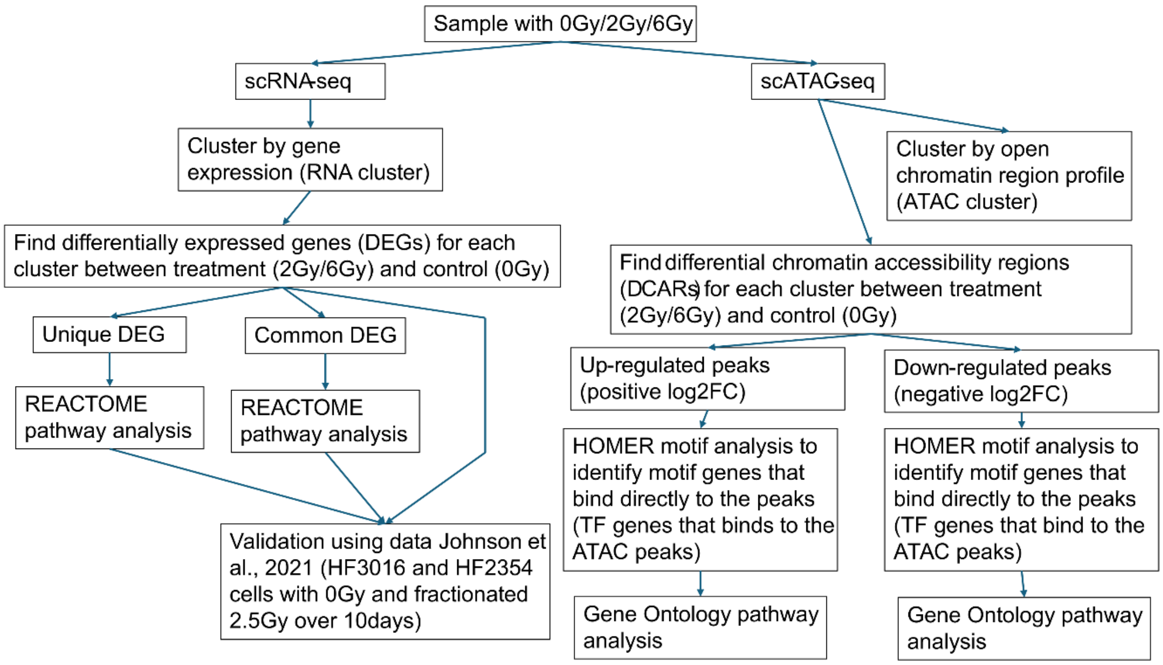


**Figure S3 Experimental workflow**

**Table S1 Pairwise comparisons of cell proportions per cluster for scRNA-seq**

| seurat_clusters | SampleID | n | ratio | total_n | p_2Gy | p_6Gy | significance |
|---|---|---|---|---|---|---|---|
| 0 | 0Gy | 138 | 0.293617021 | 470 | 0.443194545 | 0.218807978 | None |
| 0 | 2Gy | 310 | 0.31504065 | 984 | 0.443194545 | 0.218807978 | None |
| 0 | 6Gy | 151 | 0.333333333 | 453 | 0.443194545 | 0.218807978 | None |
| 1 | 0Gy | 140 | 0.29787234 | 470 | 0.106143759 | 0.660093073 | None |
| 1 | 2Gy | 252 | 0.256097561 | 984 | 0.106143759 | 0.660093073 | None |
| 1 | 6Gy | 128 | 0.282560706 | 453 | 0.106143759 | 0.660093073 | None |
| 2 | 0Gy | 70 | 0.14893617 | 470 | 0.125450843 | 0.382873327 | None |
| 2 | 2Gy | 180 | 0.182926829 | 984 | 0.125450843 | 0.382873327 | None |
| 2 | 6Gy | 78 | 0.17218543 | 453 | 0.125450843 | 0.382873327 | None |
| 3 | 0Gy | 66 | 0.140425532 | 470 | 0.684803799 | 0.215497064 | None |
| 3 | 2Gy | 129 | 0.131097561 | 984 | 0.684803799 | 0.215497064 | None |
| 3 | 6Gy | 78 | 0.17218543 | 453 | 0.684803799 | 0.215497064 | None |
| 4 | 0Gy | 56 | 0.119148936 | 470 | 0.878834793 | 1.55869E-05 | 6Gy |
| 4 | 2Gy | 113 | 0.114837398 | 984 | 0.878834793 | 1.55869E-05 | 6Gy |
| 4 | 6Gy | 18 | 0.039735099 | 453 | 0.878834793 | 1.55869E-05 | 6Gy |

**Table S2 Pairwise comparisons of GBM subtype proportions per cluster scRNA-seq**

| Seurat clusters | SampleID | predicted state | n | ratio | total_n | p_2Gy | p_6Gy | significance |
|---|---|---|---|---|---|---|---|---|
| 0 | 0Gy | Classical | 20 | 0.042553191 | 470 | 4.4498E-12 | 3.08887E-08 | Both |
| 0 | 0Gy | Mesenchymal | 59 | 0.125531915 | 470 | 3.6143E-08 | 0.050559844 | 2Gy |
| 0 | 0Gy | Proneural | 59 | 0.125531915 | 470 | 0.086063448 | 0.205715962 | None |
| 0 | 2Gy | Classical | 173 | 0.175813008 | 984 | 4.4498E-12 | 3.08887E-08 | Both |
| 0 | 2Gy | Mesenchymal | 44 | 0.044715447 | 984 | 3.6143E-08 | 0.050559844 | 2Gy |
| 0 | 2Gy | Proneural | 93 | 0.094512195 | 984 | 0.086063448 | 0.205715962 | None |
| 0 | 6Gy | Classical | 69 | 0.152317881 | 453 | 4.4498E-12 | 3.08887E-08 | Both |
| 0 | 6Gy | Mesenchymal | 38 | 0.08388521 | 453 | 3.6143E-08 | 0.050559844 | 2Gy |
| 0 | 6Gy | Proneural | 44 | 0.097130243 | 453 | 0.086063448 | 0.205715962 | None |
| 1 | 0Gy | Classical | 29 | 0.061702128 | 470 | 1.69444E-08 | 1.10854E-08 | Both |
| 1 | 0Gy | Mesenchymal | 64 | 0.136170213 | 470 | 1.10485E-12 | 1.02098E-06 | Both |
| 1 | 0Gy | Proneural | 47 | 0.1 | 470 | 0.000478196 | 0.010575131 | Both |
| 1 | 2Gy | Classical | 169 | 0.171747967 | 984 | 1.69444E-08 | 1.10854E-08 | Both |
| 1 | 2Gy | Mesenchymal | 34 | 0.034552846 | 984 | 1.10485E-12 | 1.02098E-06 | Both |
| 1 | 2Gy | Proneural | 49 | 0.049796748 | 984 | 0.000478196 | 0.010575131 | Both |
| 1 | 6Gy | Classical | 85 | 0.187637969 | 453 | 1.69444E-08 | 1.10854E-08 | Both |
| 1 | 6Gy | Mesenchymal | 19 | 0.041942605 | 453 | 1.10485E-12 | 1.02098E-06 | Both |
| 1 | 6Gy | Proneural | 24 | 0.052980132 | 453 | 0.000478196 | 0.010575131 | Both |
| 2 | 0Gy | Classical | 34 | 0.072340426 | 470 | 0.000513097 | 0.03531289 | Both |
| 2 | 0Gy | Mesenchymal | 30 | 0.063829787 | 470 | 0.003041431 | 0.308979355 | 2Gy |
| 2 | 0Gy | Proneural | 6 | 0.012765957 | 470 | 0.6744698 | 1 | None |
| 2 | 2Gy | Classical | 134 | 0.136178862 | 984 | 0.000513097 | 0.03531289 | Both |
| 2 | 2Gy | Mesenchymal | 29 | 0.029471545 | 984 | 0.003041431 | 0.308979355 | 2Gy |
| 2 | 2Gy | Proneural | 17 | 0.017276423 | 984 | 0.6744698 | 1 | None |
| 2 | 6Gy | Classical | 52 | 0.114790287 | 453 | 0.000513097 | 0.03531289 | Both |
| 2 | 6Gy | Mesenchymal | 21 | 0.046357616 | 453 | 0.003041431 | 0.308979355 | 2Gy |
| 2 | 6Gy | Proneural | 5 | 0.011037528 | 453 | 0.6744698 | 1 | None |
| 3 | 0Gy | Classical | 14 | 0.029787234 | 470 | 0.001571969 | 0.003196421 | Both |
| 3 | 0Gy | Mesenchymal | 25 | 0.053191489 | 470 | 0.0103018 | 1 | 2Gy |
| 3 | 0Gy | Proneural | 27 | 0.057446809 | 470 | 0.034773002 | 0.441964565 | 2Gy |
| 3 | 2Gy | Classical | 72 | 0.073170732 | 984 | 0.001571969 | 0.003196421 | Both |
| 3 | 2Gy | Mesenchymal | 25 | 0.025406504 | 984 | 0.0103018 | 1 | 2Gy |
| 3 | 2Gy | Proneural | 32 | 0.032520325 | 984 | 0.034773002 | 0.441964565 | 2Gy |
| 3 | 6Gy | Classical | 34 | 0.075055188 | 453 | 0.001571969 | 0.003196421 | Both |
| 3 | 6Gy | Mesenchymal | 24 | 0.052980132 | 453 | 0.0103018 | 1 | 2Gy |
| 3 | 6Gy | Proneural | 20 | 0.04415011 | 453 | 0.034773002 | 0.441964565 | 2Gy |
| 4 | 0Gy | Classical | 8 | 0.017021277 | 470 | 0.003321963 | 0.622824472 | 2Gy |
| 4 | 0Gy | Mesenchymal | 16 | 0.034042553 | 470 | 0.048769662 | 0.016221369 | Both |
| 4 | 0Gy | Proneural | 32 | 0.068085106 | 470 | 0.140168721 | 0.000687026 | 6Gy |
| 4 | 2Gy | Classical | 50 | 0.050813008 | 984 | 0.003321963 | 0.622824472 | 2Gy |
| 4 | 2Gy | Mesenchymal | 16 | 0.016260163 | 984 | 0.048769662 | 0.016221369 | Both |

| | | | | | | | | |
|---|---|---|---|---|---|---|---|---|
| 4 | 2Gy | Proneural | 47 | 0.047764228 | 984 | 0.140168721 | 0.000687026 | 6Gy |
| 4 | 6Gy | Classical | 5 | 0.011037528 | 453 | 0.003321963 | 0.622824472 | 2Gy |
| 4 | 6Gy | Mesenchymal | 4 | 0.008830022 | 453 | 0.048769662 | 0.016221369 | Both |
| 4 | 6Gy | Proneural | 9 | 0.01986755 | 453 | 0.140168721 | 0.000687026 | 6Gy |

**Table S3 Pairwise comparisons of cell proportions per cluster for scATAC-seq**

| seurat_clusters | SampleID | n | ratio | total_n | p_2Gy | p_6Gy | significance |
|---|---|---|---|---|---|---|---|
| 0 | 0Gy | 237 | 0.504255319 | 470 | 0.837992908 | 0.065334272 | None |
| 0 | 2Gy | 489 | 0.49695122 | 984 | 0.837992908 | 0.065334272 | None |
| 0 | 6Gy | 200 | 0.441501104 | 453 | 0.837992908 | 0.065334272 | None |
| 1 | 0Gy | 196 | 0.417021277 | 470 | 0.487208624 | 0.375031539 | None |
| 1 | 2Gy | 390 | 0.396341463 | 984 | 0.487208624 | 0.375031539 | None |
| 1 | 6Gy | 203 | 0.44812362 | 453 | 0.487208624 | 0.375031539 | None |
| 2 | 0Gy | 37 | 0.078723404 | 470 | 0.11256133 | 0.12537622 | None |
| 2 | 2Gy | 105 | 0.106707317 | 984 | 0.11256133 | 0.12537622 | None |
| 2 | 6Gy | 50 | 0.110375276 | 453 | 0.11256133 | 0.12537622 | None |

**Table S4 Pairwise comparisons of RNA cluster cell proportions per ATAC clusters**

| Seurat clusters | SampleID | predicted state | n | ratio | total_n | p_2Gy | p_6Gy | significance |
|---|---|---|---|---|---|---|---|---|
| 0 | 0Gy | RNA_C0 | 67 | 0.142553191 | 470 | 0.065358916 | 0.216836668 | None |
| 0 | 0Gy | RNA_C1 | 91 | 0.193617021 | 470 | 0.06463929 | 0.139459357 | None |
| 0 | 0Gy | RNA_C2 | 20 | 0.042553191 | 470 | 0.382899137 | 0.824873026 | None |
| 0 | 0Gy | RNA_C3 | 27 | 0.057446809 | 470 | 0.073070786 | 0.995186465 | None |
| 0 | 0Gy | RNA_C4 | 32 | 0.068085106 | 470 | 0.973984769 | 0.000687026 | 6Gy |
| 0 | 2Gy | RNA_C0 | 180 | 0.182926829 | 984 | 0.065358916 | 0.216836668 | None |
| 0 | 2Gy | RNA_C1 | 151 | 0.153455285 | 984 | 0.06463929 | 0.139459357 | None |
| 0 | 2Gy | RNA_C2 | 54 | 0.054878049 | 984 | 0.382899137 | 0.824873026 | None |
| 0 | 2Gy | RNA_C3 | 35 | 0.035569106 | 984 | 0.073070786 | 0.995186465 | None |
| 0 | 2Gy | RNA_C4 | 69 | 0.070121951 | 984 | 0.973984769 | 0.000687026 | 6Gy |
| 0 | 6Gy | RNA_C0 | 79 | 0.174392936 | 453 | 0.065358916 | 0.216836668 | None |
| 0 | 6Gy | RNA_C1 | 70 | 0.154525386 | 453 | 0.06463929 | 0.139459357 | None |
| 0 | 6Gy | RNA_C2 | 17 | 0.037527594 | 453 | 0.382899137 | 0.824873026 | None |
| 0 | 6Gy | RNA_C3 | 25 | 0.055187638 | 453 | 0.073070786 | 0.995186465 | None |
| 0 | 6Gy | RNA_C4 | 9 | 0.01986755 | 453 | 0.973984769 | 0.000687026 | 6Gy |
| 1 | 0Gy | RNA_C0 | 60 | 0.127659574 | 470 | 0.182690133 | 0.752291574 | None |
| 1 | 0Gy | RNA_C1 | 37 | 0.078723404 | 470 | 0.530562424 | 0.448782416 | None |
| 1 | 0Gy | RNA_C2 | 48 | 0.10212766 | 470 | 0.907586879 | 0.933980189 | None |
| 1 | 0Gy | RNA_C3 | 30 | 0.063829787 | 470 | 0.206869771 | 0.130252558 | None |
| 1 | 0Gy | RNA_C4 | 21 | 0.044680851 | 470 | 0.484689925 | 0.030480947 | 6Gy |
| 1 | 2Gy | RNA_C0 | 101 | 0.102642276 | 984 | 0.182690133 | 0.752291574 | None |
| 1 | 2Gy | RNA_C1 | 67 | 0.068089431 | 984 | 0.530562424 | 0.448782416 | None |
| 1 | 2Gy | RNA_C2 | 104 | 0.105691057 | 984 | 0.907586879 | 0.933980189 | None |
| 1 | 2Gy | RNA_C3 | 83 | 0.084349593 | 984 | 0.206869771 | 0.130252558 | None |
| 1 | 2Gy | RNA_C4 | 35 | 0.035569106 | 984 | 0.484689925 | 0.030480947 | 6Gy |
| 1 | 6Gy | RNA_C0 | 62 | 0.136865342 | 453 | 0.182690133 | 0.752291574 | None |
| 1 | 6Gy | RNA_C1 | 43 | 0.094922737 | 453 | 0.530562424 | 0.448782416 | None |
| 1 | 6Gy | RNA_C2 | 48 | 0.105960265 | 453 | 0.907586879 | 0.933980189 | None |
| 1 | 6Gy | RNA_C3 | 42 | 0.092715232 | 453 | 0.206869771 | 0.130252558 | None |
| 1 | 6Gy | RNA_C4 | 8 | 0.017660044 | 453 | 0.484689925 | 0.030480947 | 6Gy |
| 2 | 0Gy | RNA_C0 | 11 | 0.023404255 | 470 | 0.624022563 | 1 | None |
| 2 | 0Gy | RNA_C1 | 12 | 0.025531915 | 470 | 0.447849491 | 0.625638938 | None |
| 2 | 0Gy | RNA_C2 | 2 | 0.004255319 | 470 | 0.020674673 | 0.007459504 | Both |
| 2 | 0Gy | RNA_C3 | 9 | 0.019148936 | 470 | 0.327236444 | 0.7570189 | None |
| 2 | 0Gy | RNA_C4 | 3 | 0.006382979 | 470 | 0.814315292 | 0.642469197 | None |
| 2 | 2Gy | RNA_C0 | 29 | 0.029471545 | 984 | 0.624022563 | 1 | None |
| 2 | 2Gy | RNA_C1 | 34 | 0.034552846 | 984 | 0.447849491 | 0.625638938 | None |
| 2 | 2Gy | RNA_C2 | 22 | 0.022357724 | 984 | 0.020674673 | 0.007459504 | Both |
| 2 | 2Gy | RNA_C3 | 11 | 0.011178862 | 984 | 0.327236444 | 0.7570189 | None |
| 2 | 2Gy | RNA_C4 | 9 | 0.009146341 | 984 | 0.814315292 | 0.642469197 | None |
| 2 | 6Gy | RNA_C0 | 10 | 0.022075055 | 453 | 0.624022563 | 1 | None |
| 2 | 6Gy | RNA_C1 | 15 | 0.033112583 | 453 | 0.447849491 | 0.625638938 | None |
| 2 | 6Gy | RNA_C2 | 13 | 0.028697572 | 453 | 0.020674673 | 0.007459504 | Both |
| 2 | 6Gy | RNA_C3 | 11 | 0.024282561 | 453 | 0.327236444 | 0.7570189 | None |
| 2 | 6Gy | RNA_C4 | 1 | 0.002207506 | 453 | 0.814315292 | 0.642469197 | None |
